# The *T. ispahanicum* elongated glume locus *P2* maps to chromosome 6A and is associated with the ectopic expression of *SVP-A1*

**DOI:** 10.1101/2022.01.27.478079

**Authors:** Yi Chen, Yinqi Liu, Junli Zhang, Adam Torrance, Nobuyoshi Watanabe, Nikolai M. Adamski, Cristobal Uauy

## Abstract

In rice and wheat, glume and floral organ length are positively correlated with grain size, making them an important target to increase grain size and potentially yield. The wheat subspecies *Triticum ispahanicum* is known to develop elongated glumes and floral organs as well as long grains. These multiple phenotypic effects are controlled by the *P2* locus, which was previously mapped to wheat chromosome 7B. Using three mapping populations, we show that the long glume locus *P2* does not map to chromosome 7B, but instead maps to a 1.68 Mbp interval on chromosome 6A. Within this interval, we identified *SVP-A1*, a MADS box transcription factor which is the direct ortholog of the maize gene underlying the ‘pod corn’ *Tunicate* locus and is a paralog to the *T. polonicum* elongated glume *P1* gene. In *T. ispahanicum*, we identified a private allele which has a 482-bp deletion in the *SVP-A1* promoter and is associated with ectopic and higher expression of *SVP-A1* in the elongated glumes and floral organs. We used near-isogenic lines (NILs) to show that *P2* has a consistent positive effect on the length of glume, lemma, palea, spike and grain. Based on the mapping data, natural variation, biological function of *SVP* genes in cereals and expression analyses, we propose the MADS-box transcription factor *SVP-A1* as a promising candidate for *P2*.

**Key message:** We propose the MADS-box transcription factor *SVP-A1* as a promising candidate gene for the elongated glume locus *P2*, which maps to chromosome 6A instead of the previously proposed chromosome 7B.

## Introduction

Inflorescence architecture influences final grain yield in crops, including cereals such as wheat (*Triticum aestivum*), rice (*Oryza sativa*), and maize (*Zea mays*). In cereals, the inflorescence is composed of specialized floret-bearing branches known as spikelets, which are subtended by sterile bract-like organs called glumes. Each floret is composed of two leaf-like sheathing structures, the lemma and the palea, as well as lodicules, stamens, and the pistil. Increasing the number of spikelets (Wolde et al. 2019), the number of fertile florets (*GNI1*; Sakuma et al. 2019), and the size/weight of the grain can increase the final grain yield in cereals, including wheat (Feng et al. 2018). Grain size was shown to be positively correlated in rice and wheat with the size of floral organs, including lemma and palea (Lombardo and Yoshida 2015; Millet 1986; reviewed in Li and Li 2015). This is likely because the lemma and palea envelop the developing grain in wheat and rice and thus define the space that the grain can grow into. These observations suggest that we can modify grain size and weight through manipulation of floral organ size in wheat and rice.

Several subspecies of wheat have elongated glumes and floral organs in comparison to hexaploid bread wheat (*T. aestivum*). These wheat subspecies represent valuable genetic resources with the potential to improve floral organ size and potentially grain size in wheat. Tetraploid subspecies (4x) *T. polonicum* and *T. ispahanicum*, and hexaploid subspecies (6x) *T. petropavlovsky* were originally identified and classified as subspecies due to their long-glume phenotype (Khoshbakht 2009; Wang et al. 2002; Watanabe 1999; Watanabe and Imamura 2002; Watanabe et al. 1998), which is often accompanied by elongated lemmas and paleae. Mimicking their long glume and floral organ phenotypes, these wheat subspecies also produce long and slender grains (Gegas et al. 2010). Identification of the genes underlying the elongated glume and floral organ phenotypes in these subspecies could therefore provide gene targets to increase grain length in wheat.

The gene underlying the long-glume phenotype of *T. polonicum* and *T. petropavlovsky* was previously mapped to the *P1* locus on chromosome 7A (Watanabe et al. 1996). Recently, several groups have independently discovered that the long-glume and lemma phenotypes of *P1* are caused by the ectopic expression of *VEGETATIVE TO REPRODUCTIVE TRANSITION 2* (*VRT2*), a MADS-box transcription factor belonging to the *SHORT VEGETATIVE PHASE* (*SVP*)/*StMADS11-like* subfamily (Adamski et al. 2021; Liu et al. 2021; Schilling et al. 2020; Xiao et al. 2021). Interestingly, genes from this subfamily also influence glume and lemma length across different grass species. Ectopic expression of *ZMM19* leads to the elongated glume phenotype observed in ‘pod corn’ maize (Han et al. 2012; Wingen et al. 2012), ectopic expression of *BM1* in barley leads to elongated lemmas and paleae (Trevaskis et al. 2007) and ectopic expression of *OsMADS22* leads to elongated glumes in rice (Sentoku et al. 2005). Based on these examples, the expression of *SVP/StMADS11-like* genes seems to be linked with the size of glumes and floral organs in cereals.

The causal gene for the long-glume phenotype in *T. ispahanicum* has yet to be identified. *T. ispahanicum* was characterized in the Isfahan province in Iran in the middle of the 20^th^ century by independent expeditions (Heslot 1959; Kihara et al. 1965; Kuckuck 1956). The cultivation of *T. ispahanicum* has since disappeared likely due to its susceptibility to different diseases (Khoshbakht 2009). The initial grouping of *T. ispahanicum* as a subspecies was based solely on its long-glume characteristic. Since then, karyotypic information and whole genome sequencing analyses have shown low genetic diversity within *T. ispahanicum* accessions and that *T. ispahanicum* is genetically similar to domesticated emmer *T. dicoccon* (Badaeva et al. 2015; Zhou et al. 2020).

Previous studies have mapped the long-glume phenotype of *T. ispahanicum* to the *P2* locus on chromosome 7B, and *P2* was therefore hypothesised to be the B-genome homoeolog of *P1* (Watanabe 1999; Watanabe et al. 2002). Here, we used multiple genetic approaches and populations to fine-map the *P2* locus to a 1.68 Mbp interval on chromosome 6A. Using near-isogenic lines, we characterized the effect of *P2* on yield-related traits, including glume length, maternal floral organ size, grain, and inflorescence morphology. Within the 1.68 Mbp physical interval, we identified *SVP-A1* (*TraesCS6A02G313800*), a *SVP*/*StMADS11-like* gene, which is the closest paralog of the *VRT2* gene underlying *P1*. Based on allelic variation, biological function of *SVP*/*StMADS11-like* genes in cereals, and expression analysis, we propose *SVP-A1* as a promising candidate for *P2*.

## Materials and Methods

### Germplasm

We made two F_2_ mapping populations to map *P2*. We obtained *T. ispahanicum* accessions, ‘T1120002’ and ‘TRI 7117’ from the John Innes Centre (JIC) Germplasm Resources Unit (GRU) and the IPK Genebank, respectively (Supplementary Figure 1). We crossed T1120002 (T1) and TRI 7117 (TRI) to *T. durum* cultivar ‘Langdon’ (LDN) to create two F_2_ populations T1 × LDN (*n* = 93 F_2_ individuals) and TRI × LDN (*n* = 120 F_2_ individuals), respectively. To confirm the mapping of *P2* to chromosome 6A, we developed a F_2_ mapping population using previously published BC_6_ near-isogenic lines (NILs), LD222 and P2-LD222 (Watanabe 1999). The NILs were developed by crossing *T. ispahanicum* ‘CL1120001’ to durum wheat cultivar ‘LD222’. Subsequent backcrossing to the recurrent parent LD222 while selecting for the elongated glume phenotype yielded P2-LD222 with the *P2* locus from *T. ispahanicum*. P2-LD222 was named ‘ANW 5B’ in Watanabe et al. (2003) but here we use the original nomenclature. We crossed the NILs to make the F_2_ population LD222 × P2-LD222 (*n* = 172 F_2_ individuals). To further map *P2* on chromosome 6A, we self-pollinated F_2_ lines that were heterozygous between 401 and 602 Mbp on chromosome 6A from both the T1 × LDN and the TRI × LDN populations. We identified 70 F_3_ heterozygous recombinant lines between markers *M1* and *M7*, of which we selected seven for self-pollination to recover homozygous recombinant lines in the F_3:4_ generation. We performed an additional round of fine-mapping by identifying 11 F_3_ heterozygous recombinant lines between markers *M4* and *M6*, which were self-pollinated to recover homozygous recombinant lines.

We used two pairs of *P2* NILs to characterise the effects of *P2* on yield-related traits in the field and glasshouse as well as to characterise expression levels of candidate genes. The first set of NILs includes the previously described LD222 and P2-LD222. The second set includes LD222(*Rht-B1b*) and P2-LD222(*Rht-B1b*). LD222(*Rht-B1b*) was described as ‘ANDW 4A’ in previous literature and is a LD222 near-isogenic line (BC_6_) with an *Rht-B1b* introgression from the durum wheat cultivar ‘Cando’ (Watanabe et al. 2003). To develop P2-LD222(*Rht-B1b*), we crossed LD222(*Rht-B1b*) with P2-LD222 and select F_2_ progenies with elongated glume and semi-dwarfism phenotypes. Homozygosity at both loci were then confirmed by phenotyping the F_3_. The first set of NILs is referred to as the LD222 NILs while the second set of NILs is referred to as LD222(*Rht-B1b*).

The recurrent parents with normal glume length phenotype and the *SVP-A1a* allele are described as *P2*^*WT*^ while the near-isogenic counterparts with elongated glumes and the *P2* introgression (including *SVP-A1b* from *T. ispahanicum*) are described as *P2*^*ISP*^.

For allelic diversity studies, we used accessions of *T. ispahanicum, T. polonicum, T. petropavlovskyi, T. dicoccoides, T. turgidum* L. ssp. *durum*, and *T. aestivum* from the IPK Genebank, the USDA-ARS National Small Grains Collection (NSGC), and the JIC GRU.

### Glasshouse and field phenotyping

For the initial mapping in the T1 × LDN and TRI × LDN F_2_ populations, we grew plants in the glasshouse under long day conditions (16-h light: 8-h dark) and measured the glume length of the two central spikelets of the primary spike for each F_2_ progeny using a ruler (*n* = 93 T1 × LDN F_2_ individuals; *n* = 120 TRI × LDN F_2_ individuals). For all the experiments conducted in the glasshouse, the plants were grown in 1 L pots in “John Innes Cereal Mix” (65% peat, 25% loam Soil, 10% grit, 3 kg/m^3^ dolomitic limestone, 1.3 kg/m^3^ PG mix and 3 kg/m^3^ osmocote exact).

For each round of fine-mapping, seven (recombinants between markers *M1* and *M7*; Supplementary Table 1) and eleven (recombinants between markers *M4* and *M6*; Supplementary Table 2) F_3:4_ homozygous recombinant lines, alongside their non-recombinant sibling line, were phenotyped for glume length of the two central spikelets of the primary spike (four glumes per plant, *n* = 5 plants per genotype). We also phenotyped the main spike of individual plants for spike length and spikelet number. Spike length was measured as the distance between the lowest rachis node to the tip of the terminal spikelet. Spikelet number was counted as the total number of spikelets per spike, regardless of fertility.

For characterisation of the *P2* NILs, we grew the LD222 NIL pair in 1 L pots in the glasshouse under long day conditions (16-h light: 8-h dark). We dissected the main spike from each plant (*n* = 3 plants per genotype) to measure the size of glume, lemma, palea, and grain across the entire spike as described in Adamski et al. (2021).

Both the LD222 and LD222(*Rht-B1b*) *P2* NILs were evaluated at the John Innes Centre Experimental Field Station in Bawburgh, UK (52°37’50.7”N 1°10’39.7”E). The *P2* NILs were sown in the spring of 2021 in 1 m^2^ plots in a randomized complete block design with six blocks. We collected five representative primary spikes (subsamples) from each genotype per each of the six blocks to measure spike length, spikelet number, and number of fertile florets. The number of fertile florets per spikelet was estimated based on the number of grains produced by the two central spikelets. We measured the length of glume, lemma and palea by measuring these tissues from the first floret of the four central spikelets with a ruler. Grain morphological traits were measured using the MARVIN grain analyser (GTA Sensorik GmbH, Neubrandenburg, Germany) and grains per spike were counted from these spikes. We measured heading days as the number of days from sowing to reach the day when at least 75% of primary spikes within the plot were fully emerged. Plant height was measured as the distance from the soil to the tip of the spike excluding awns.

### Grain developmental time course

To investigate the effect of *P2* on grain development, we collected five primary spikes per genotype (LD222 and LD222(*Rht-B1b*) NILs) per block at 0, 5, 10, 14, and 19 days post-anthesis (dpa). Due to the elongated glume, anthesis was difficult to detect based on anther extrusion alone. We therefore defined anthesis as the time that the anthers of the central spikelet turned yellow. For each spike, we collected four developing ovaries/grains from florets 1 and 2 of the two central spikelets (4 ovaries/grains × 5 spikes = 20 ovary/grain sample per block). Ovary and grain morphology was measured using the MARVIN grain analyser.

### Genotyping

For the initial mapping in the F_2_ populations, we identified polymorphisms for marker development by aligning RNA-sequencing data of *T. ispahanicum* (BioProject PRJNA288606; Zou et al. 2015) and exome-capture sequence data of Langdon (BioProject PRJNA684023; Adamski et al. 2021) against a “tetraploid” RefSeqv1.0 (IWGSC et al. 2018) that lacked the D-genome. We called single nucleotide polymorphisms (SNPs) with Freebayes (v1.1.0) using standard filters and minimum alternate count of 5 (Garrison and Marth 2012). Based on the predicted SNPs, we developed 53 Kompetitive Allele-Specific PCR (KASP) markers across the 14 chromosomes using PolyMarker (Ramirez-Gonzalez et al. 2015) (Supplementary Table 3). The cycle conditions for the KASP assay were: 15 mins at 94 °C; 10 touchdown cycles of 20 seconds at 94 °C followed by 65 °C to 57 °C for 1 min; 40 cycles of 94 °C for 20s, 57 °C for 1 min. This program was used for all the KASP markers described in this study.

To check the isogenic status of the LD222 NILs and to identify polymorphisms for marker development, we genotyped LD222 and P2-LD222 using the Breeders’ 35K Axiom Array (Allen et al. 2017). We only kept markers from the A- and B-genome that belonged to the “PolyHighRes”, “NoMinorHom”, and “OTV” cluster categories (9879 markers in total; Supplementary Table 4). After removing monomorphic markers, we were left with 133 markers. For F_2_ mapping in the LD222 × P2-LD222 population, we converted these polymorphisms on chromosomes 2A, 2B, and 6A into KASP markers using PolyMarker (Ramirez-Gonzalez et al. 2015) (Supplementary Table 3).

### Statistical analysis

F_2_ QTL mapping was performed using the R/qtl package version 1.5 in R studio using the single-QTL genome scan with normal model and the EM algorithm (Broman et al. 2003). In addition, we performed analysis of variance (ANOVA) to test the effect of key markers on chromosomes 6A, 7A and 7B on glume length and performed Tukey’s test to compare each genotypic group. For the F_3:4_ homozygous recombinant lines, we performed *t*-tests between the glume length values of the recombinant plants against their non-recombinant siblings. To characterize the association of *P2* with quantitative traits (spike length and spikelet number) that have large variation between F_3:4_ homozygous recombinant families, we performed ANOVA based on haplotype group. We grouped homozygous recombinants based on their haplotype from markers *M11* to *M14*. Lines with *T. ispahanicum* allele from *M11* to *M14* were grouped into the *T. ispahanicum* haplotype while lines with the Langdon allele from *M11* to *M14* were assigned to the wildtype group. We performed two-way ANOVA for the T1 × LDN and TRI × LDN populations separately to evaluate the effect of the *P2* haplotype group (accounting for recombinant family) on spike length and spikelet number. Planned contrasts were performed to test if haplotype groups were significantly different (Supplementary Table 5).

To assess the effect of *P2* on glume and floral organ morphology across the spike, we classified the spike into three distinct regions that were independent of spikelet number. The apical region of the spike contained the data from the apical 25% of the spikelets, the central region of the spike contained the data from the middle 50% of spikelets, while the basal region of the spike contained the basal 25% of spikelets. We performed two-way ANOVA to test the effect of the *P2* allele, position on the spike (basal, central, apical) and their interaction on the size of glume, lemma, grain and palea (Supplementary Table 6).

For the grain developmental time course, we performed ANOVA to test the effect of *P2*, timepoint and their interaction (accounting for block effect) on grain morphometrics in each pair of NILs separately. We used the average of 20 grains measured per biological replicate as input for the analysis. For pericarp cell length analysis, we performed two-way ANOVA to test the effect of *P2*, pericarp cell position on the grain and their interaction. For field-based data, we performed two-way ANOVA to test the effect of *P2*, the NIL background and their interaction (accounting for block effect) on the traits measured. For all the phenotypes, we used the average of five spikes from each block (subsamples) for the analysis. Following every ANOVA, planned contrasts were performed to test if *P2*^*WT*^ *and P2*^*ISP*^ were significantly different.

### Candidate gene identification

To characterize the genes within the *P2* mapping interval, we used BioMart (Kinsella et al. 2011) to extract the RefSeqv1.1 gene model annotation (IWGSC et al. 2018), GO annotation, and the closest orthologs in *O. sativa* and *Arabidopsis thaliana*. We used the funRiceGenes database (https://funricegenes.github.io/) to investigate the function of these genes in rice (Yao et al. 2017).

### Phylogenetic analysis of *SVP*/*StMADS11-like* genes in grasses

We identified the closest orthologs of the three wheat *SVP*/*StMADS11-like* genes in *T. durum, Brachypodium distachyon, Hordeum vulgare, O. sativa* and *Z. mays* based on Plant Compara from Ensembl Plant (Howe et al. 2020; Schilling et al. 2020). We aligned their amino acid sequences in MEGA X using MUSCLE with default settings (Kumar et al. 2018). From the alignment, we generated a phylogeny tree using the Maximum likelihood method with bootstrap method and 1000 bootstrap replication under default setting in MEGA X.

### Allelic variation analyses

We characterized the sequence variation of *VRT-B2* (*TraesCS7B02G080300*) and *SVP-A1* (*TraesCS6A02G313800*) in *T. ispahanicum* including 1500 bp up- and downstream of the untranslated region of the genes. We used the whole genome sequencing reads of *T. ispahanicum* accessions from Zhou et al. (2020), including ‘KU-145’, KU-4580’, ‘PI 294477’, ‘PI 284478’, ‘PI 354293’, ‘PI 330548’, and ‘TRI 6177’. The sequencing reads were aligned to a ‘tetraploid’ version of the RefSeqv1.0 assembly (IWGSC et al. 2018) lacking the D-genome chromosomes using HiSat2-v-2.1.0 with default settings (Kim et al. 2019). The alignments were visualized using Integrated Genomics Viewer (IGV; Robinson et al. 2011) and polymorphisms of *VRT-B2* and *SVP-A1* were called based on visual inspection of the alignment. We only recorded sequence variations with a read depth of at least three reads and which were present in at least five out of the seven *T. ispahanicum* accessions (to allow for low sequence coverage in samples). The 482-bp promoter deletion of *SVP-A1* was then confirmed via Sanger sequencing of PCR amplicons. For Sanger sequencing, we performed PCR amplification (95°C for 3 mins; 35 cycles of 95°C for 15 s, 59°C for 45 s, 72°C for 90 s; 2 mins of 72°C) on genomic DNA extracted from wheat seedlings using the PCR promoter deletion markers (Supplementary Table 7).

To characterize the sequence of *SVP-A1* and *VRT-B2* in *T. dicoccon*, which has normal glume size and was proposed as the progenitor of *T. ispahanicum* (Badaeva et al. 2015), we used the whole genome sequencing reads of five *T. dicoccon* accessions from Zhou et al. (2020). This included ‘PI 532305’, ‘PI 266842’, ‘CItr 3686’, ‘PI 626391’, and ‘PI 94668’. We used the same approach as described above and recorded sequence variations that were present in at least three of the five *T. dicoccon* accessions. We also used the alignments of *T. ispahanicum* accessions against the Chinese Spring RefSeqv1.0 reference to investigate the allelic status of the *BTR-A1 (*3A: 65,869,056-65,869,644*), BTR-B1* (3B:88,971,298-88,977,068), and *Q* (*TraesCS5A02G473800*) loci. These same sequences were also compared against wild emmer wheat *T. dicoccoides* (Zavitan, WEWSeq_v1.0; Avni et al. 2017) and domesticated durum wheat *T. turgidum* L. ssp. *durum* (Svevo.v1; Maccaferri et al. 2019).

Lastly, we screened diverse tetraploid and hexaploid wheat accessions for the *SVP-A1* promoter deletion and the missense mutation c.431A>G that led to exon five p.Q144R substitution (nomenclature based on Den Dunnen and Antonarakis 2001). These accessions were screened using either a KASP assay, Sanger sequencing, or analysis of whole-genome sequencing data from Zhou et al. (2020) as described previously (Supplementary Table 8). This included 181 *T. aestivum*, 396 *T. turgidum ssp. durum*, 13 *T. ispahanicum*, 5 *T. dicoccon* and 11 *T. dicoccoides* accessions. We also screened a subset of accessions collected during the Kuckuck expedition to Iran (Kuckuck 1956) including 63 *T. aestivum*, 1 *T. spelta*, 29 *T. turgidum ssp. durum* and 5 *T. dicoccon* accessions (Supplementary Table 9).

### Pericarp cell length measurement

To compare pericarp cell length of *P2*^*WT*^ and *P2*^*ISP*^, we imaged pericarp surfaces of LD222 NILs grown in the glasshouse using scanning electron microscopy (SEM). We collected two grains from the first floret of the two central spikelets from five independent plants grown in separate 1 L pots. Dry grain samples were mounted crease-down onto 12.5 mm SEM specimen stubs (Agar Scientific Ltd). We sputter coated each sample with 7.5 nm gold using a high vacuum sputter coater (Leica EM ACE600; Leica Microsystem). We imaged the grain surface at 3 kV with the Nova NanoSEM450 (FEI, United States) at the top (brush side), middle, and bottom (germ side) of the grain with one image in each section (Fig 4). Pericarp cell length then was measured using Fiji (Schindelin et al. 2012) and we calculated the median cell length for each image.

### RNA collection

The LD222 NILs were grown in 1 L pots in the glasshouse under long day conditions (16h light: 8 h dark). We harvested flag leaf, as well as glume, lemma, palea, and anthers at Waddington stage 7.5-8 (Waddington et al. 1983) from central spikelets of the main spike of four plants for each NIL (four biological replicates). We also collected grains from florets one and two at 3, 10, and 20 dpa from the four central spikelets (eight grains per sample) of the main spike of four plants for each NIL (four biological replicates). All tissues were immediately frozen in liquid nitrogen and stored at −80 °C.

Grains were homogenized using mortar and pestle with liquid nitrogen while other tissues were homogenized in SPEX CertiPrep 2010-230 Geno/Grinder (Cat No.: 12605297, Fischer Scientific) using 5 mm steel beads (Cat No.: 69989, Qiagen). For grain samples, RNA was extracted following the protocol described in Adamski et al. (2021). For non-grain tissues, we used the Spectrum Plant Total RNA kit (Cat No.: STRN250-1KT, Sigma) following the manufacturer’s protocol.

### Reverse transcription quantitative PCR (RT-qPCR)

We performed reverse transcription using the SuperScript III First-Strand Synthesis System (Cat No.: 18080051, Thermo Fisher). One microgram of RNA was used as input and the reaction was performed with Oligo(dT) primer following the manufacturer’s protocol. We used the LightCycler 480 SYBR Green I Master Mix (Roche Applied Science, UK) to perform RT-qPCR in a LightCycler 480 II instrument (Roche Applied Science, UK). The cycle conditions were: 5 min at 95 °C; 45 cycles of 10 s at 95 °C, 15 s at 62 °C, 30 s at 72 °C; dissociation curve from 60 °C to 95 °C to determine primer specificity. Reactions were performed using three technical replicates per sample. Relative gene expression was calculated using the 2^-ΔΔCT^ method (Livak and Schmittgen 2001) with *Actin* as the reference gene and a common calibrator to produce relative expression values that were comparable across samples.

### Phylogenetic shadowing and MEME motif discovery

We defined 2000 bp upstream of the transcription start site as the putative promoter sequence of *SVP-A1* (*TraesCS6A02G313800*). We then performed a phylogenetic shadowing analysis with mVista (Frazer et al. 2004) for the *SVP-A1* promoter and the orthologous sequences from barley (*H. vulgare*; *HORVU6Hr1G077300*), *B. distachyon* (*BRADI3g58220*), rice (*Os02g0761000*), maize (*GRMZM2G370777*), and sorghum (*Sorghum bicolor*; *SORBI_3004G306500*). We searched for conserved regions in 20 bp windows with a minimum length of 15 bp and a minimum sequence identity of 85%.

Given that mVISTA requires positional conservation of motifs, we used the ‘MEME’ tool of MEME Suite 5.3.3 to discover motifs that are only conserved in sequence (Bailey et al. 2009). Predicted motifs were parsed through the ‘Tomtom’ tool of MEME Suite 5.3.3 to compare them against known motifs from the ‘JASPAR Core non-redundant plant motif’ database (Fornes et al. 2020).

Lastly, we obtained the promoter sequence of *ZMM19* from ‘pod corn’ (Wingen et al. 2012) and aligned it against the wildtype *ZMM19* (*GRMZM2G370777*) allele to determine the presence of the three identified motifs with respect to the ‘pod corn’ promoter re-arrangement.

## Results

### The *P2* locus of *T. ispahanicum* maps to chromosome 6A and not to chromosome 7B as previously proposed

Previous studies had proposed that *P2* was homoeologous to *P1* (Watanabe 1999). Recently, multiple groups identified *VRT-A2* as the causal gene for *P1* (Adamski et al. 2021; Liu et al. 2021; Xiao et al. 2021). We therefore compared the sequence of *VRT-B2* (*TraesCS7B02G080300*) in five *T. ispahanicum* accessions to Chinese Spring (RefSeqv1.0). We identified 33 polymorphisms with 32 in non-coding regions and one silent mutation on exon seven (Supplementary Table 10). We also investigated the allelic diversity of *VRT-B2* in *T. dicoccon* accessions (*n* = 5), which have normal glume size (∼10 mm) and are proposed to be the progenitor of *T. ispahanicum*. All polymorphisms identified in the *T. ispahanicum VRT-B2* gene were present in the *T. dicoccon* accessions and are therefore not private to *T. ispahanicum*. This suggests that these polymorphisms are not the causal events underlying the elongated glume of *T. ispahanicum* (Supplementary Table 10) and that *VRT-B2* is unlikely to underlie *P2*.

To further define the *P2* locus on chromosome 7B, we developed two F_2_ mapping populations between Langdon (normal glume size; ∼10 mm) and two separate *T. ispahanicum* accessions, T1120002 (T1) and TRI 7117 (TRI) (Supplementary Figure 1). These *T. ispahanicum* accessions are expected to carry *P2* given their elongated glume phenotype (glume size ∼17 mm and ∼15 mm, respectively). We designed twelve markers in 50-100 Mbp intervals along chromosome 7B and used them to genotype both mapping populations (Supplementary Table 3). Based on the glume length phenotype of the two F_2_ populations (T1 × LDN, *n* = 93 F_2_ plants; TRI × LDN, *n* = 120 F_2_ plants), we performed QTL analysis. Surprisingly, we found no significant association between the markers on chromosome 7B and glume length in either population (Fig 1; Supplementary Table 3; Supplementary Figure 2). This includes KASP markers at 62 Mbp (*7B:62046061*) and 105 Mbp (*7B:105882162*), which flank *VRT-B2* (90.1 Mbp). Our F_2_ mapping results are therefore not consistent with previously published maps that position *P2* on chromosome 7B (Watanabe 1999).

**Fig 1.**
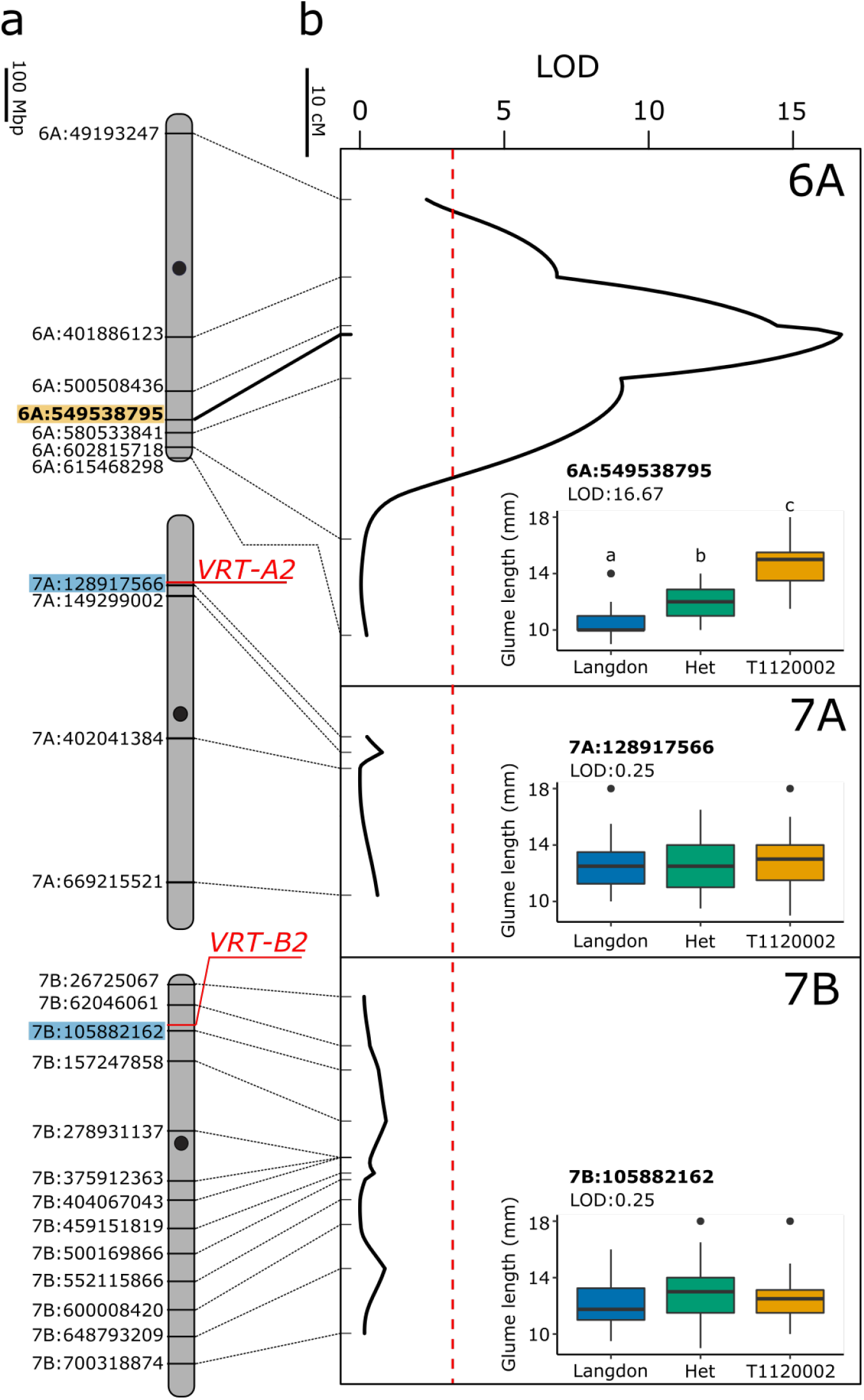
The long glume trait of *T. ispahanicum* maps to chromosome 6A in the T1 × LDN F_2_ population. **a**, Physical maps of chromosomes 6A, 7A and 7B as well as the KASP markers used for mapping. Approximate centromere positions are denoted by the black circle. The location of *VRT2* homoeologs is highlighted in red, with the closest KASP markers in blue. The peak marker on chromosome 6A is highlighted in orange. **b**, QTL analysis of glume length for chromosomes 6A, 7A and 7B. Genetic positions of markers are denoted with ticks and are connected to their physical position in (a). The dashed red line denotes the significance threshold (LOD > 3.0). **Inset** Glume length distribution of F_2_ individuals (*n* = 93) carrying parental or heterozygous genotypes at markers *6A:549538795, 7A:128917566*, and *7B:105882162*. Significant differences between genotypic classes were only found for the 6A marker using ANOVA and post-hoc Tukey test (*P* < 0.01)

To map the elongated glume trait of these two *T. ispahanicum* accessions, we developed KASP markers across all chromosomes and genotyped the T1 × LDN population. We found significant associations between the markers on chromosome 6A and glume size, with the peak marker at 549.5 Mbp (*6A:549539795*; LOD = 16.7; Fig 1b, Supplementary Figure 2, Supplementary Table 11). Lines homozygous for the *T. ispahanicum* allele at this marker had longer glumes (14.7 mm) than lines homozygous for the Langdon allele (10.7 mm; *P* <0.01). Lines that were heterozygous at this allele developed an intermediate glume size (12.1 mm), significantly different to both homozygous allele classes, suggesting a semi-dominant effect of *P2*. We did not find an association between glume length and markers on other chromosomes including marker *7A:128917566*, which is less than 150 kbp distal of *VRT-A2* (*TraesCS7A02G175200*; 128.7 Mbp), the casual gene for *P1* (Fig 1; Supplementary Figure 2). Consistent with the result in T1 × LDN, we found a significant association between markers on chromosome 6A and glume length with the same peak marker, *6A:549539795* (LOD = 13.2), in the TRI × LDN F_2_ population (Supplementary Figure 2b). Together, these genetic results suggest that the elongated glume locus in the two *T. ispahanicum* accessions maps to chromosome 6A.

Our results contradicted previous publications that placed the elongated glume locus of *T. ispahanicum* onto chromosome 7B. The original germplasm stock of *T. ispahanicum* used to map *P2* (accession CL1120001; Watanabe 1999) originated from the same germplasm repository as the conspicuously named accession T1120002 used in this study. We could not, however, discard the possibility that the two *T. ispahanicum* accessions used in the current study carried a previously uncharacterised locus for elongated glumes different to *P2*. We therefore acquired the original materials that were used to map *P2* onto chromosome 7B; the recurrent parent LD222 with normal glume size and its near-isogenic sibling P2-LD222, which carries the *T. ispahanicum P2* introgression and develops elongated glumes (Watanabe 1999). We used the 35K Axiom array (Allen et al., 2017) to genotype the NILs and found that intervals on chromosomes 2A, 2B, and 6A were polymorphic between the NILs (Supplementary Figure 3). However, there were few or no detectable polymorphisms on other chromosomes including chromosome 7B (no polymorphic markers out of 721 genotyped markers), which suggests that most chromosomes, apart from 2A, 2B, and 6A, are monomorphic between the NILs.

To test if *P2* maps to chromosomes that are polymorphic between the NILs, we created a F_2_ mapping population by crossing the NILs, LD222 × P2-LD222 (*n* = 172 F_2_ plants). We designed markers on chromosome 2A, 2B, and 6A based on the 35K Axiom array probes. Consistent with our results from the other F_2_ mapping populations (T1 × LDN and TRI × LDN), we found a significant association between glume length and markers on chromosome 6A (Supplementary Figure 2b). We did not find a significant association between markers on chromosome 2A and glume length. Markers on chromosome 2B appeared to be monomorphic in the F_2_ population (Supplementary Figure 4). We speculate that this is due to similarity of the 2B array probes to homoeologous regions on 2A. These genomic and genetic results in the original germplasm used to map *P2*, alongside the mapping data of two independent *T. ispahanicum* mapping populations suggest that the *P2* elongated glume locus is located on chromosome 6A and not on chromosome 7B as previously described.

### Fine-mapping of the *P2* locus to a 1.68 Mbp region on chromosome 6A

To fine-map the *P2* locus, we selected F_2_ lines that were heterozygous on chromosome 6A from 401 Mbp to 602 Mbp from both the T1 × LDN and TRI × LDN F_2_ populations. We screened over 2000 F_3_ plants and identified 70 heterozygous recombinants on chromosome 6A within the 401-602 Mbp interval. Initially, we phenotyped glume length in seven F_3:4_ homozygous recombinant lines between markers *M1* and *M7* (and their non-recombinant siblings) and mapped the *P2* locus to a 21 Mbp interval between markers *M4* (541 Mbp) and *M6* (562 Mbp; Fig 2a; Supplementary Table 1; Supplementary Table 12). To further define the *P2* locus, we developed ten additional KASP markers and advanced eleven F_3:4_ homozygous recombinant lines with recombination events between markers *M4* and *M6* from both populations. We determined their *P2* status based on the glume length phenotype and pairwise comparisons against their corresponding non-recombinant sibling line. The key recombination events R4 and R8 delimited *P2* to a 1.68 Mbp region on chromosome 6A between markers *M11* (549,037,133 bp) and *M14* (550,717,813 bp; Fig 2b, c; Supplementary Table 2).

**Fig 2.**
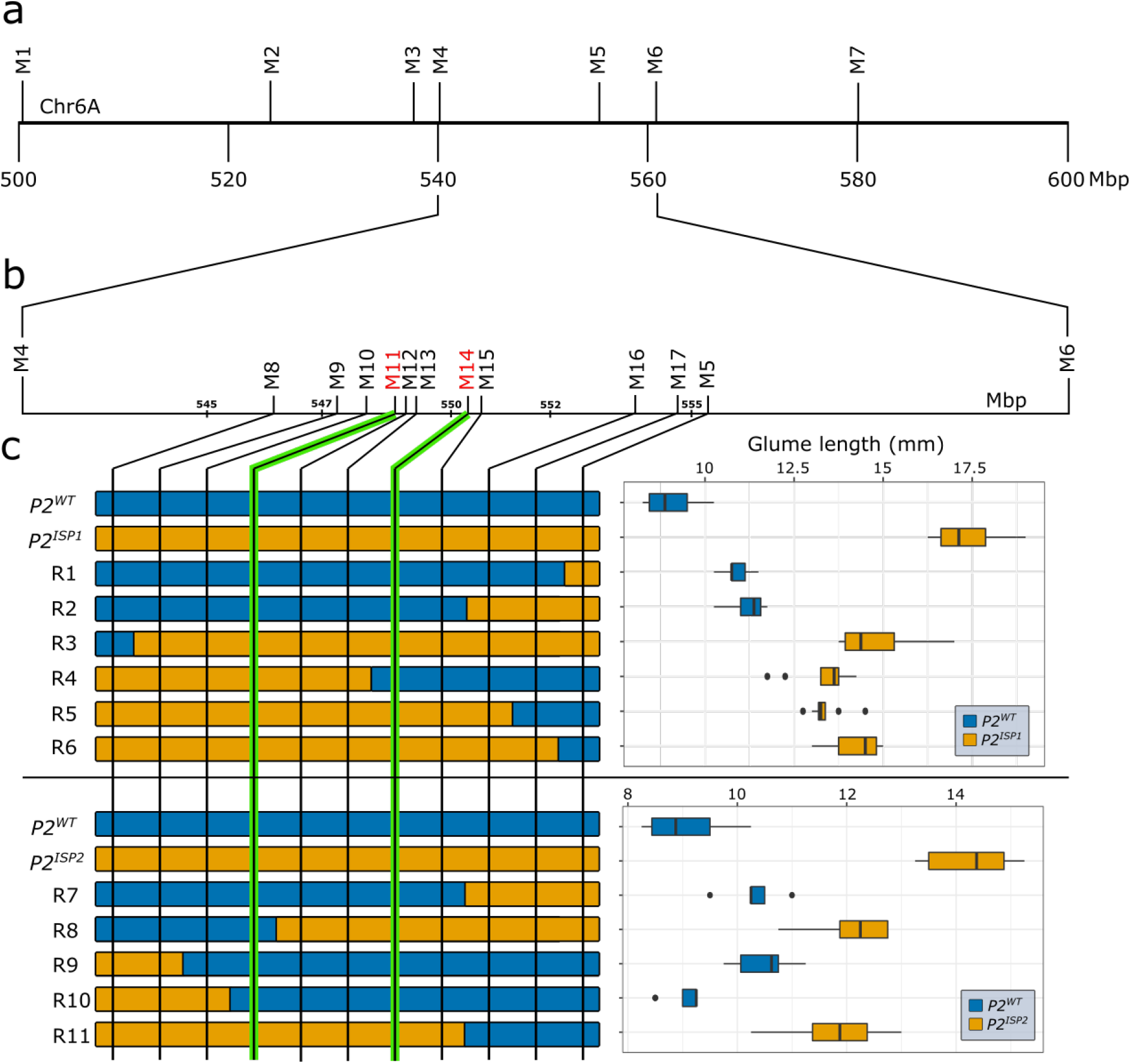
Fine mapping of the *P2* locus using F_3:4_ homozygous recombinant lines from T1 × LDN and TRI × LDN mapping populations. **a**, The glume length phenotype of *P2* was initially mapped between markers *M4* and *M6* (∼21 Mbp) using seven F_3:4_ recombinant lines (Supplementary Table 1). **b**, The interval was further delineated to a 1.68 Mbp interval between markers *M11* and *M14* using the glume length phenotype from 11 F_3:4_ recombinant lines (Supplementary Table 2). **c**, Graphical genotype of eleven critical recombinants between markers *M5* and *M8* from the cross of Langdon to either T1120002 (*ISP1*, top panel) or TRI 7117 (*ISP2*, bottom panel). Each F_3:4_ homozygous recombinant line was determined to carry *P2*^*WT*^ or *P2*^*ISP*^ based on the glume length phenotype (*n* = 8 plants) and pairwise comparison against its non-recombinant sibling line (Supplementary Table 2). The *P2* interval is defined by recombinant lines R4 and R8 between markers *M11* and *M14* (highlighted with green lines). The box plots show the middle 50% of the data with the median represented by the vertical line. Whisker represents datapoint within 1.5 times the interquartile range with outliers highlighted as individual dots

We observed higher spikelet number and longer spike length in the F_3:4_ recombinants with the elongated glume phenotype. However, given the more quantitative nature of these traits, we were unable to map them to a single major locus using the homozygous recombinant lines. We therefore tested whether the *P2* region was associated with variation in spikelet number and spike length, by grouping recombinant lines based on their 1.68 Mbp haplotype from markers *M11* to *M14*. We found that the *T. ispahanicum P2* haplotype group was associated with significantly longer spikes (*P* < 0.001) and higher spikelet number (*P* < 0.01) in both populations (Supplementary Table 5). Our results show that the 1.68 Mbp *T. ispahanicum P2* locus is associated with longer glumes and spikes, and higher spikelet number, compared to the Langdon wildtype.

### *SVP-A1* is a promising candidate gene for *P2*

Using the RefSeqv1.1 annotation, we identified 15 high-confidence (HC) and 20 low-confidence (LC) genes within the 1.68 Mbp *P2* interval (Supplementary Table 13). The HC genes have Gene Ontology (GO) functional annotation including GTPase, cellulase, ubiquitin-protein transferase, and hydrolase. Among the 15 HC genes, we identified *SVP-A1* (*TraesCS6A02G313800*; GO: 0030154 flower development), a MADS-box gene belonging to the *SHORT VEGETATIVE PHASE (SVP)*/*StMADS11-like* subfamily. We conducted a phylogenetic analysis of *SVP*/*StMADS11-like* proteins from representative grasses including *T. aestivum, T. durum, B. distachyon, H. vulgare, O. sativa*, and *Z. mays* (Supplementary Figure 5). We identified three clades consistent with a monocot-specific triplication of *StMADS11-like* genes. Proteins from each clade have been shown to influence glume or lemma length when ectopically expressed in wheat (Adamski et al. 2021; Liu et al. 2021; Xiao et al. 2021), maize (Han et al. 2012; Wingen et al. 2012), rice (Sentoku et al. 2005), and barley (Trevaskis et al. 2007). The biological function of previously characterized *SVP*/*StMADS11-like* protein in grasses, alongside the fine-mapping data, makes *SVP-A1* a promising candidate gene for *P2*.

### The *T. ispahanicum SVP-A1* allele, which is private to this subspecies, includes a 482-bp promoter deletion and an A431G missense mutation

We characterized the sequence of *SVP-A1* in *T. ispahanicum* (*n* = 7 accessions) by comparing their genome sequencing data (Zhou et al. 2020) to the Chinese Spring *SVP-A1* sequence. We identified a 482-bp deletion in the promoter region (455 bp upstream of the ATG start codon), 54 SNPs/small indels in the non-coding region, and one missense mutation on exon five (6A:g.550640120T>C that leads to c.431A>G and p.Q144R substitution; Supplementary Table 14). The 54 SNPs/small indels were present in multiple *T. dicoccon* accessions that had normal glume length, suggesting they are unlikely to influence glume length. The 482-bp promoter deletion is of interest as previous studies have established a link between *SVP*/*StMADS11-like* gene expression and glume length (Adamski et al. 2021; Wingen et al. 2012). The A431G missense mutation led to an amino acid substitution (Q144R) within the K-box domain of the MADS-box transcription factor, a domain which is required for hetero- and homo-dimer formation (Riechmann et al. 1996). The glutamine at position 144 is conserved across the *SVP1* and *VRT2* clade except for *ZMM21*, whereas the *SVP3* clade proteins have a lysine residue at this position (Supplementary Figure 6). Therefore, the Q144R amino acid substitution could potentially affect *SVP-A1* function. We thus focused on the 482-bp promoter deletion and the A431G missense mutation, which were not found in the *T. dicoccon* genome sequences but were present in all 13 *T. ispahanicum* accessions examined. We denoted the *SVP-A1* allele from *T. ispahanicum*, including the 482-bp promoter deletion and A431G polymorphism, as *SVP-A1b* (Figure 3A).

**Fig 3.**
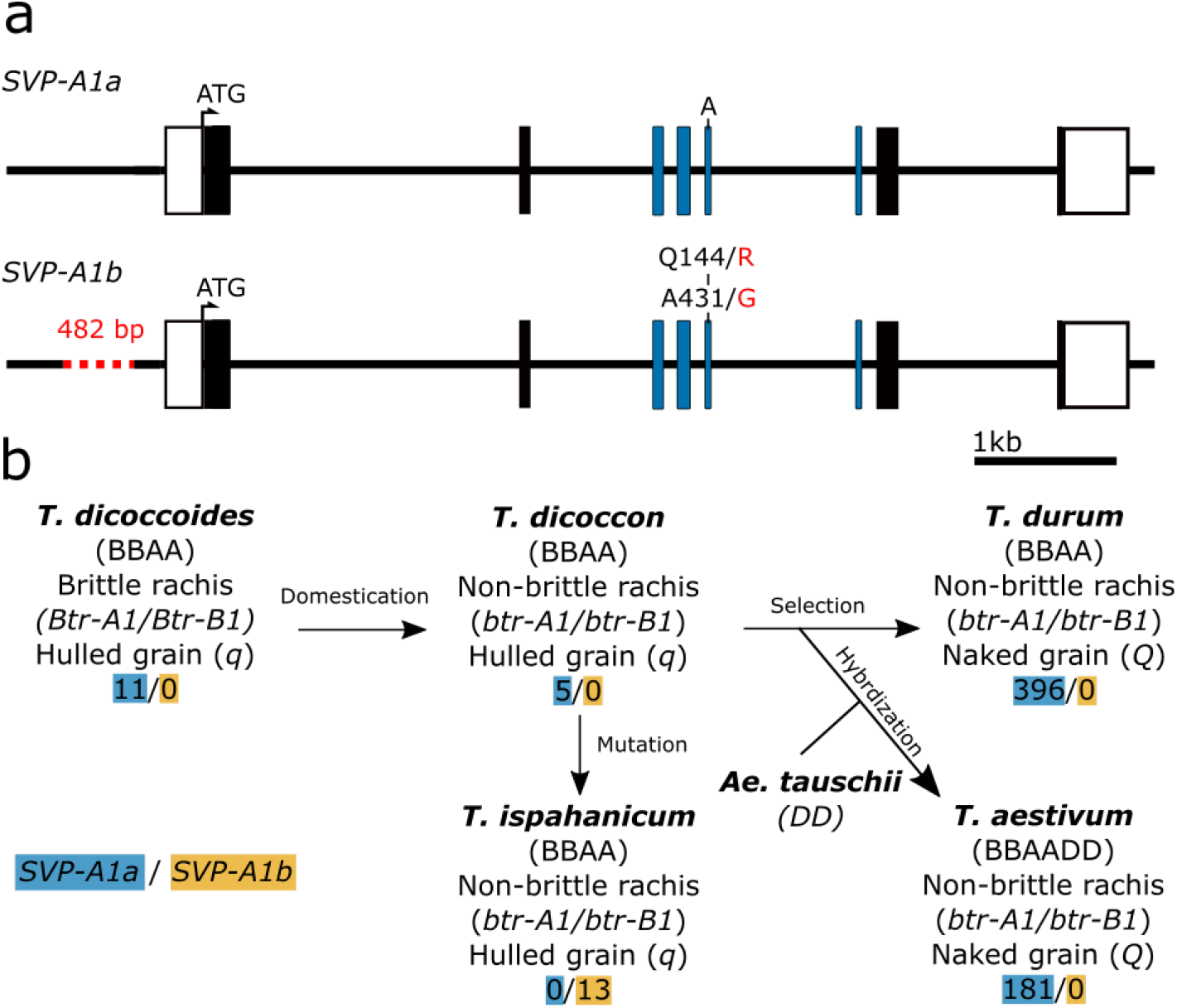
*SVP-A1* allelic diversity and prevalence of the *T. ispahanicum SVP-A1b* allele in tetraploid and hexaploid wheat germplasm. **a**, The *T. ispahanicum SVP-A1b* allele has a 482-bp promoter deletion and a Q144R amino acid substitution with respect to the Chinese Spring wildtype *SVP-A1a* allele. White boxes denote UTR and black/coloured boxes are exons. The exons encoding for the K-box domain are depicted in blue. **b**, Simplified diagram of the evolution and domestication of tetraploid and hexaploid wheat with the proposed origin of *T. ispahanicum*. The number of accessions that carry the wildtype *SVP-A1a* allele (blue) or the *SVP-A1b* allele with the 482-bp promoter deletion and A431G missense mutation that led to Q144R substitution (orange) is shown. All wheat accessions shown have normal sized glumes, apart from the 13 *T. ispahanicum* accessions, which have elongated glumes. The allelic status of *Non-brittle rachis 1* (*Btr-1*) homoeologs and the hulled grain *q* gene are shown

**Fig 4.**
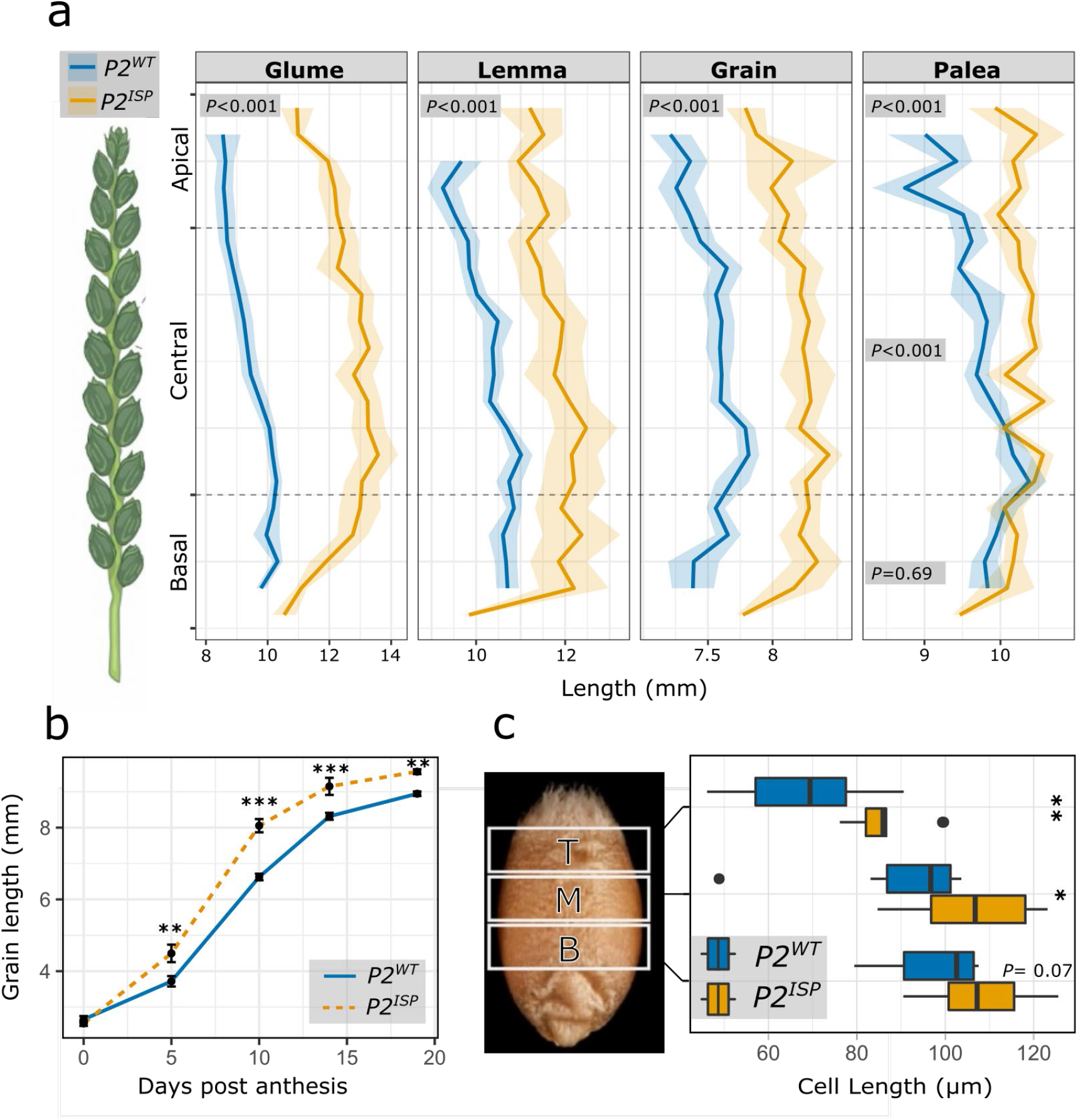
Phenotypic effect of *P2* on glume, floral organ, and grain morphology in LD222 NILs. **a**, Drawing of a wheat spike and the mean glume, lemma, grain, and palea length along the spike of *P2*^*WT*^ and *P2*^*ISP*^ based on samples grown in the glasshouse (*n* = 3 plants). The apical region contains the data from the apical 25% of the spikelets, central region contains the data from middle 50% while the basal region contains the basal 25% of the spikelets. Ribbons represent the standard error. *P*-values were based on planned contrast between *P2*^*WT*^ and *P2*^*ISP*^ at the specific region of the spike (Supplementary Table 6); *P <* 0.001 for all tissues/regions except for the palea in the basal region (*P* = 0.69). **b**, Time course of ovary/grain length within the first floret of four central spikelets in field-grown *P2*^*WT*^ and *P2*^*ISP*^ NILs (*n* = 20 ovaries/grains sampled from 5 spikes per each of 6 blocks). Error bar denotes the standard error. *P*-values were based on planned contrast between *P2*^*WT*^ and *P2*^*ISP*^ at the specific timepoint **c**, Pericarp cell size comparison between glasshouse-grown *P2*^*WT*^ and *P2*^*ISP*^ NILs based on grains collected from the first floret of two central spikelet (*n* = 10 grains from 5 plants). The grain image depicts the three regions in which an image was taken including top (T), middle (M), and bottom (B). The box shows the middle 50% of the data with the median represented by the vertical line. Whiskers represent datapoints within 1.5 times the interquartile range with outliers highlighted as individual dots. *P*-values were based on planned contrast between *P2*^*WT*^ and *P2*^*ISP*^ at the specific region of the grain. \**P* < 0.05; \*\**P* < 0.01; \*\*\**P* < 0.001

To investigate the prevalence of the *SVP-A1b* allele in wheat, we screened global germplasm for the 482-bp promoter deletion and the A431G polymorphism using KASP genotyping, Sanger sequencing, and analysis of available whole genome sequencing data. Across 185 hexaploid and 422 tetraploid wheat accessions, we did not find any accession that carried either the 482-bp promoter deletion or the A431G polymorphism. In contrast, both polymorphisms were present in all 13 *T. ispahanicum* accessions examined (Fig 3b; Supplementary Table 8). This screen included other long-glume wheat subspecies, such as *T. polonicum* (*n* = 10) and *T. petropavlovskyi* (*n* = 4), for which *VRT-A2* is known to be the underlying genetic locus (Adamski et al. 2021; Liu et al. 2021). The Kuckuck expedition, which collected the first *T. ispahanicum* accessions, also collected additional wheat accessions from regions in Iran (Kuckuck 1956). Since these wheat accessions were growing in the same space and time as *T. ispahanicum*, we hypothesized that these accessions might also carry *SVP-A1b*. We genotyped a subset of accessions from the Kuckuck expedition (63 *T. aestivum*, 1 *T. spelta*, 29 *T. turgidum ssp. durum* and 5 *T. dicoccon*; Supplementary Table 8; Supplementary Figure 7) for *SVP-A1b* and also assessed their glume phenotype. We found that all accessions developed normal-sized glumes (5.0 -10.5 mm) and that no accession carried *SVP-A1b*. Together, these allelic variation studies suggest that the the *SVP-A1b* allele, including the 482-bp promoter deletion and A431G polymorphism, is completely linked with the long-glume phenotype of *P2* and is likely private to *T. ispahanicum*.

### *T. ispahanicum* is likely an accession of *T. dicoccon*

Given that *T. ispahanicum* was proposed to originate from domesticated emmer (*T. dicoccon*) (Badaeva et al. 2015, Khoshbakht 2009, Zhou et al. 2020), we investigated whether *T. ispahanicum* accessions carry the same alleles that confer non-brittle rachis and hulled grain in *T. dicoccon* (Avni et al. 2017; Sang 2009). We confirmed through *in-silico* analysis that *T. ispahanicum* has the non-brittle rachis alleles (*btr-A1* and *btr-B1*) on chromosomes 3A and 3B, respectively, and the hulled grain allele, *qq*, which are defining characteristics of *T. dicoccon*. The similarity in chromosome structure and the low sequence variation between *T. ispahanicum* and *T. dicoccon* suggest that *T. ispahanicum* was derived from *T. dicoccon* likely of West Asia origin. We therefore propose *T. ispahanicum* should not be considered as a subspecies but rather as an accession of *T. dicoccon*.

### The *P2* locus influences the size of glumes, maternal floral organs, and grain morphology

To characterize the effect of *P2* on various traits, we used two pair of *P2* NILs in the genetic background of LD222 carrying either the wildtype *Rht-B1a* allele or the semi-dwarfism *Rht-B1b* allele. In both NIL pairs, we confirmed that the line with the *P2* introgression had elongated glumes and carried the *SVP-A1b* allele. We performed an initial characterization of the LD222 NILs in a glasshouse experiment and with both NIL pairs (LD222 and LD222(*Rht-B1b*)) in the field.

We investigated the effects of *P2* on glume, lemma, and palea length by comparing the LD222 NILs. We observed that the *P2*^*ISP*^ NILs had significantly longer glumes and lemmas than the *P2*^*WT*^ NILs across the entire spike (Fig 4a; Supplementary Table 6). We observed significantly longer paleae in *P2*^*ISP*^ at the apical and central position but not at the basal position. Consistent with the glasshouse data for LD222, the field results with both LD222 and LD222(*Rht-B1b*) NILs showed that *P2*^*ISP*^ was associated with significantly longer glumes, lemmas, and paleae compared to *P2*^*WT*^ (Supplementary Figure 8). Our results demonstrate that the *T. ispahanicum P2* locus increases glume, lemma, and palea length independent of the *Rht-B1* allelic status.

Next, we investigated the effect of *P2* on grain morphology. Based on the glasshouse sample, we observed that *P2*^*ISP*^ was associated with longer grains across the spike (Fig 4a). *P2*^*ISP*^ was also associated with a decrease in grain width across the spike with smaller effect on the apical region (*P* < 0.05; Supplementary Table 6). Therefore, the increase in grain length associated with *P2*^*ISP*^ only contributed to an increase in grain area in the apical position of the spike (*P* = 0.03). In the field trial, we found that *P2*^*ISP*^ was associated with a significant increase in grain length in both NIL pairs (Supplementary Figure 8; Supplementary Table 15). Consistent with the glasshouse data, we observed that the grain length effect did not translate into increased grain area in LD222 NILs, because *P2*^*ISP*^ was associated with a decrease in grain width. In contrast, we did not see a significant decrease in grain width associated with *P2*^*ISP*^ in the LD222(*Rht-B1b*) NILs, resulting in larger grain area. *P2*^*ISP*^ only decreased thousand grain weight (TGW) in the LD222 NILs but not LD222(*Rht-B1b*) (Supplementary Figure 8). Our results suggest that *P2*^*ISP*^ has a consistent positive influence on grain length. However, the effect of *P2* on other grain morphometrics and TGW can be influenced by the genetic background including the *Rht-B1* allele.

We investigated the effect of *P2* on heading time, plant height, and spike morphology. As expected of the effect of *Rht-B1b*, we observed that LD222(*Rht-B1b*) NILs are shorter than LD222 NILs. However, the *P2* allele did not influence heading time nor plant height in either NIL pair under field conditions (Supplementary Figure 9). Consistent with the results of the F_3:4_ families, we found that *P2*^*ISP*^ was associated with significantly longer spikes and increased spikelet number in comparison to *P2*^*WT*^ for both NIL pairs (Supplementary Table 16). However, despite the increase in spikelet number, *P2*^*ISP*^ spikes had a significant decrease in grain number per spike due to a decrease in the number of fertile florets per spikelet (Supplementary Figure 10). Our results show that *P2* ^*ISP*^ has a consistent positive effect on spikelet number and spike length, but decreases grain number per spike.

### *P2* influences early grain development and pericarp cell length

To investigate at what developmental stage *P2* influences grain length, we sampled ovaries/grains from the LD222 and LD222(*Rht-B1b*) NILs grown in the field (*n* = 6 blocks) at 0, 5, 10, 14, and 19 days post-anthesis (dpa). We found that at 0 dpa, ovary length was similar between *P2*^*ISP*^ and *P2*^*WT*^ in both NIL pairs (Fig 4; Supplementary Table 16). However, in both NIL pairs the grain length of *P2*^*ISP*^ NILs became significantly longer than *P2*^*WT*^ at 5 dpa and remained longer at later timepoints (Supplementary Figure 11; Supplementary Table 16). This suggests that the effect of *P2*^*ISP*^ on grain length is not a pre-anthesis effect, but rather caused by increased grain elongation at the early stages of grain development. To gain an insight into the potential mechanism, we compared the pericarp cell length of *P2*^*ISP*^ and *P2*^*WT*^ in LD222 NILs using SEM. We found that *P2*^*ISP*^ NILs had significantly longer cells at the top and the middle of the grain than *P2*^*WT*^ NILs (Fig 4c; Supplementary Table 17). These results show that the *P2* mediated increase in grain length is due, at least in part, to an increase in pericarp cell length.

### *SVP-A1* is expressed ectopically in the elongated glumes, lemmas, and paleae of *P2* ^*ISP*^ NILs

We investigated the expression of *SVP* genes in flag leaves, glumes, lemmas, paleae, and anthers of LD222 *P2* NILs at Waddington stage 7.5-8. Using RT-qPCR, we did not detect expression of *SVP-A1* in *P2*^*WT*^ glumes, lemmas, and paleae (Fig 5a). In contrast, we found significant ectopic expression of *SVP-A1* in glumes, lemmas, and paleae of *P2*^*ISP*^ NILs (*P* < 0.05). The gene was expressed at similar levels in flag leaves and anthers of the two genotypes. We also investigated the expression of the B-genome homoeolog, *SVP-B1*, and found no difference in expression between the NILs in the five tissues. Similarly, we did not detect significant differences in expression of the *SVP* paralogs *VRT-A2* and *VRT-B2* when comparing *P2* NILs (Fig 5a). These results show that the *SVP-A1b* allele in the *P2*^*ISP*^ NIL is associated with ectopic expression of *SVP-A1* in the tissues that were elongated in *P2*^*ISP*^ including glume, lemma, and palea, but does not affect the expression of the B-genome *SVP1* homoeolog, nor the expression of *VRT2*.

**Fig 5.**
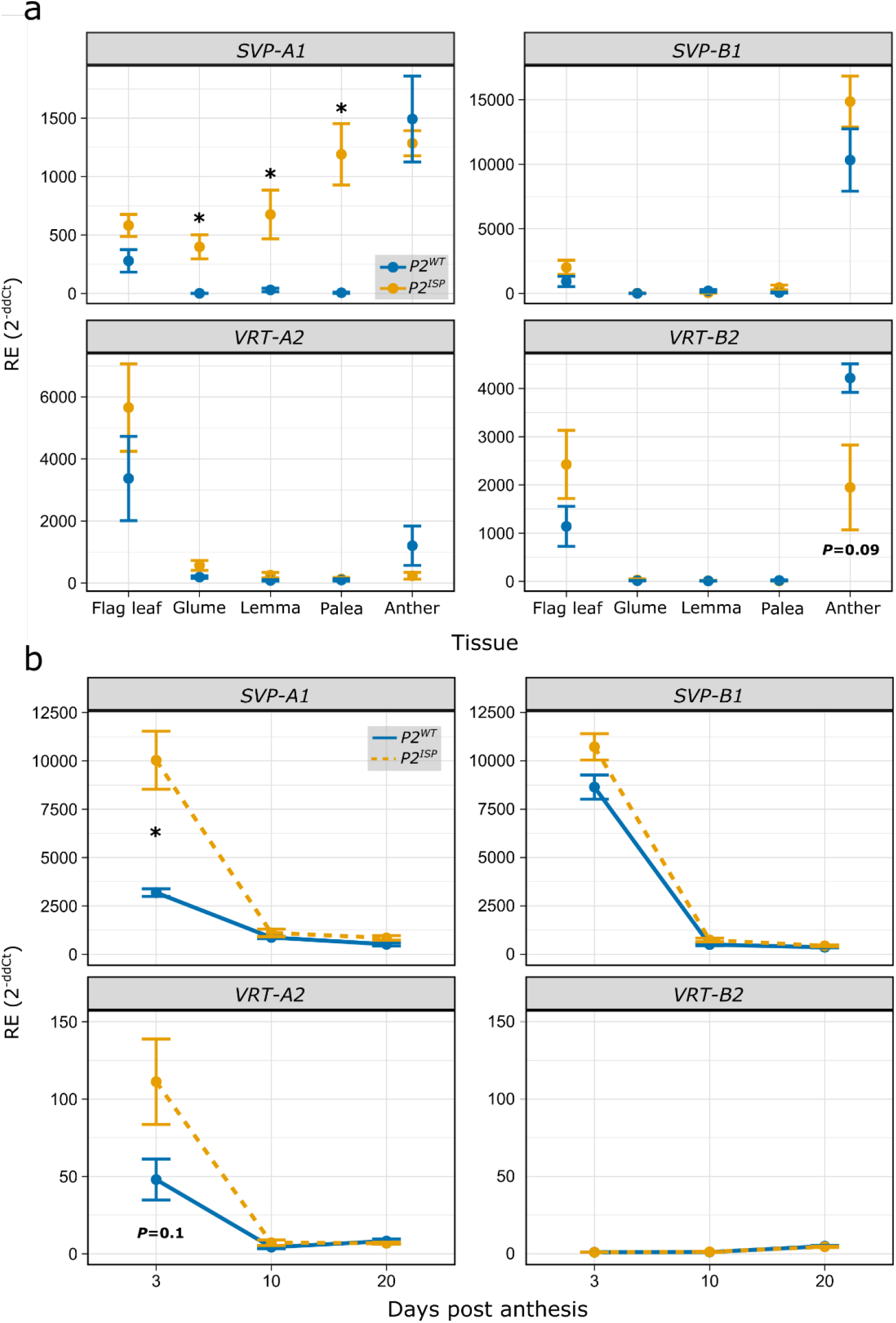
*SVP-A1* is expressed more highly and ectopically in *P2*^*ISP*^ relative to *P2*^*WT*^ NILs. **a**, Relative expression of *SVP-A1* (*TraesCS6A02G313800*), *SVP-B1* (*TraesCS6B02G343900*), *VRT-A2* (*TraesCS7A02G175200*), and *VRT-B2* (*TraesCS6B02G080300*) in flag leaf, glume, lemma, palea, and anther of LD222 *P2* NILs (*n* = 4 plants). Tissues were collected between Waddington stage 7.5 and 8. Samples were collected from floret one and two of the four central spikelets. **b**, Relative expression of *SVP-A1, SVP-B1, VRT-A2*, and *VRT-B2* in grains at 3, 10, and 20 days post-anthesis. Pairwise *t-* tests were conducted to compare relative expression levels between *P2*^*ISP*^ and *P2*^*WT*^ at each time point. Relative expression (RE) values in (a) and (b) are an average 2^ddCt^ ± standard error of the mean from four independent biological replicates per tissue/timepoint, run in triplicates. Error bars are mean ± SEM. \**P* < 0.05

Given that *P2*^*ISP*^ was associated with increased grain length during early grain development, we next investigated *SVP-A1* expression in the developing grains of LD222 *P2* NILs at 3, 10, and 20 dpa. We found that *SVP-A1* expression decreases during grain development in *P2*^*WT*^ (Fig 5b). A similar down-regulation of *SVP-A1* was found in *P2* ^*ISP*^, albeit *SVP-A1* was more highly expressed in *P2* ^*ISP*^ compared to *P2*^*WT*^ NILs at 3 dpa (but not at 10 dpa or 20 dpa). As before, we did not detect differences in expression for *SVP-B1* or *VRT2* homoeologs between *P2* alleles. Our results show an increase in *SVP-A1* expression during early grain development in the *P2* ^*ISP*^ NILs which have elongated grains.

### The *SVP-A1* promoter contains conserved sequence motifs across grasses

We showed that the *P2*^*ISP*^ NILs with the *SVP-A1b* allele (482-bp promoter deletion and A431G missense mutation) had higher and ectopic expression of *SVP-A1* in the tissues with elongated organ size (glume, lemma, palea, and grain). We therefore hypothesized that the deleted promoter region of *SVP-A1b* may contain regulatory motifs that affect its expression profile. Using phylogenetic shadowing (mVISTA, Frazer et al. 2004) of the 2 kbp upstream sequence of several grass species, we identified two motifs that are conserved in sequence and position in *SVP1* grass orthologs (Fig 6a). A third motif was identified using the MEME suite, which does not require the motifs to be positionally conserved (Fig 6b). The three motifs were located within the 482-bp deleted region of *SVP-A1b* and were conserved in their order across the grass species considered. In ‘pod corn’ maize, the duplication and promoter re-arrangement of *ZMM19* (direct wheat ortholog of *SVP-A1*) leads to its ectopic expression and the characteristic elongated glume phenotype (Han et al. 2012; Wingen et al. 2012). We found that all three motifs were also lost in maize during the promoter re-arrangement of *ZMM19*, which occurs 131 bp downstream from the end of motif three (Fig 6c). We compared the three motifs against the JASPAR Core non-redundant plant motif database and found no significant match for motif 1 and motif 2. Motif 3, however, was similar to several MADS box transcription factor binding sites (*q* < 0.05, eg: MA1203.1) as it contained a putative CArG-box. Given the sequence conservation across ∼60 million years of Poaceae divergence time, it is tempting to speculate that these motifs regulate the expression of *SVP-A1* and that their absence in *SVP-A1b* contributes to the ectopic expression of *SVP-A1* in *P2*^*ISP*^, analogous to the ectopic expression of *ZMM19* in ‘pod corn’.

**Fig 6.**
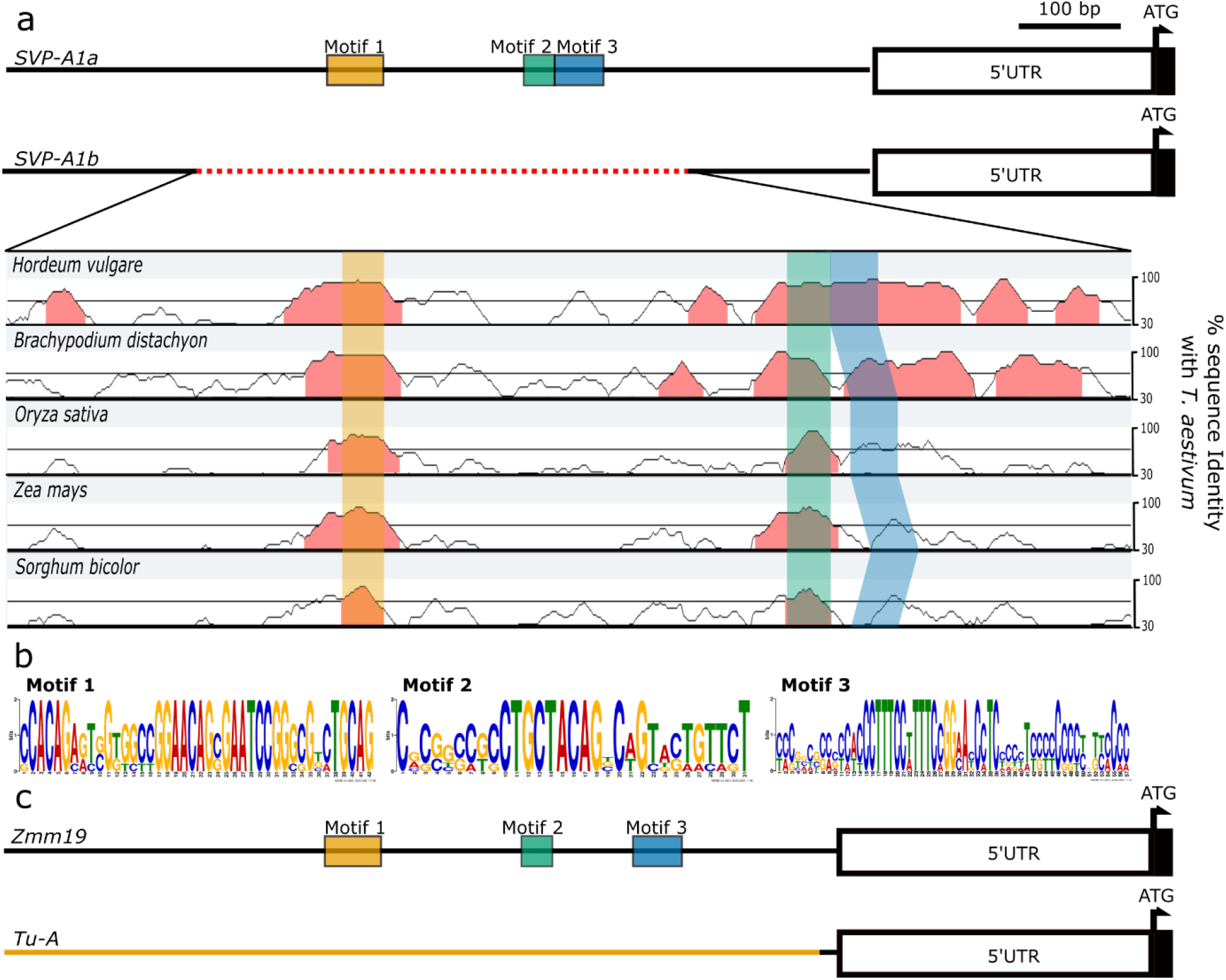
Sequence alignment of the promoter region of *SVP-A1* orthologs from several grass species identified three conserved motifs within the deleted region of *SVP-A1b*. **a**, The promoter sequence of *SVP-A1* was aligned against the closest orthologs in other grass species including *H. vulgare* (HORVU6Hr1G077300), *B. distachyon* (Bradi3G58220), *O. sativa* (Os02G0761000), *Z. mays* (GRMZM2G370777) and *S. bicolor* (SORBI_3004G306500). We used MEME suite (Bailey et al. 2015) and mVISTA (Frazer et al. 2004) to identify motifs that are conserved only in sequence or conserved in both sequence and position, respectively, along the promoters. Conserved regions (>80% similarity over a 20-bp sliding window) are highlighted in red. Motif 1 and motif 2 were identified using mVISTA while motif 3 was identified by MEME. **b**, Consensus logo of the three motifs identified. **c**, *Tunicate* (*Tu-A*) in maize has elongated glumes due to a promoter rearrangement (highlighted in orange, based on sequencing information from Wingen et al. (2012)) that leads to the loss of the three conserved Poaceae *SVP1* motifs.

## Discussion

### *P2* is located on chromosome 6A

A previous study (Watanabe 1999) assigned the *T. ispahanicum P2* long-glume locus to chromosome 7B. Based on this, our initial hypothesis was that *P2* represented the B-genome homoeolog of *VRT-A2*, the *T. polonicum P1* locus for elongated glumes on chromosome 7A. However, in two F_2_ mapping populations (T1 × LDN, TRI × LDN) using different accessions of *T. ispahanicum* as parents, we could not detect a significant association between glume length and any of the markers on chromosome 7B, including markers flanking *VRT-B2* (Fig 1b; Supplementary Figure 2b). This was supported by the lack of unique allelic variation in *VRT-B2* across *T. ispahanicum* accessions. Instead, the QTL mapping showed significant associations between glume length and chromosome 6A markers, with *6A:549538795* as the common peak marker in both F_2_ populations. These results suggest that *VRT-B2* does not underlie *P2*, and that the major effect locus from both *T. ispahanicum* accessions used here is located on chromosome 6A.

For the original mapping of *P2* to chromosome 7B, Watanabe (1999) used phenotypic linkage between the long-glume phenotype of *T. ispahanicum* accession CL1120001 and two other phenotypes (seedling and culm colour) controlled by genes on chromosome 7B. To test whether accession CL1120001 carried a different long-glume locus than the two *T. ispahanicum* accessions used here, we genotyped the original *P2* near-isogenic lines (P2-LD222 and its recurrent parent LD222) used by Watanabe (1999). We found detectable polymorphisms between the NILs only on chromosomes 2A, 2B, and 6A (Supplementary Figure 3). Furthermore, we generated a F_2_ mapping population between the NILs and observed a significant association between glume length and marker *6A:549538795* on chromosome 6A, but no association with chromosome 7B (Supplementary Figure 2b). Thus, based on both the original and novel germplasm and molecular markers, we conclude that *P2* maps to chromosome 6A. It is worth noting that, except for chromosome 2A, 6A, 7A, and 7B, we designed only a limited number of markers across the other chromosomes. We therefore cannot exclude the possibility that there are additional *T. ispahanicum* QTL that influence glume length on other chromosomes.

### *SVP-A1* is a promising candidate gene for *P2*

The phenocopy on spike morphology between *P1* and *P2* (e.g., increase in glume, lemma, and spike length) suggested that the gene underlying *P2* functions via a similar mechanism or is in the same genetic pathway as *VRT-A2*. We fine-mapped the *P2* locus to a 1.68 Mbp interval on chromosome 6A, including 15 HC genes (Fig 2). These genes included *SVP-A1*, a MADS-box transcription factor of the *SVP*/*StMADS11-like* family and the closest wheat paralog to *VRT-A2*. Previous studies have shown that higher/ectopic expression of members from all three *SVP*/*StMADS11-like* clades in cereals increase the length of glume and floral organs (e.g., lemma and palea). Natural variation in glume length in *T. polonicum* and ‘pod corn’ are caused by the ectopic expression of *VRT-A2* (Adamski et al. 2021; Liu et al. 2021) and the *SVP1* maize ortholog *ZMM19* (Han et al. 2012; Wingen et al. 2012), respectively. Similarly, transgenic overexpression of *OsMADS22* in rice (Sentoku et al. 2005) and of *HvBM1* in barley (Trevaskis et al. 2007) led to elongated glume and lemma, respectively. Based on these previous studies, alongside the genetic and expression analyses presented here, we propose that the elongation of glume, lemma, and palea in *T. ispahanicum* is associated with the ectopic expression of *SVP-A1*. While transgenic experiments will be required to establish a more direct and causal link in this relationship, the proposed mechanism is reminiscent of the natural variation that leads to ectopic expression of *SVP/StMADS11-like* genes *in* ‘pod corn’ and *T. polonicum*.

Previous work has illustrated the importance of conserved non-coding sequences (CNSs) in regulating the spatial and temporal expression pattern of genes (Meng et al. 2021) across related species. For example, targeted modification of the CNSs at the promoter region of *WOX9* led to the same drastic changes in the inflorescence architecture across tomato and groundcherry, two distantly related Solanaceae species (Hendelman et al. 2021). In this study, we found that the *SVP-A1b* allele of *T. ispahanicum* carries a 482-bp promoter deletion that is private to *T. ispahanicum* and encompasses three conserved motifs across *SVP/StMADS11-like* genes in cereals (Fig 3, Fig 6a). These three motifs were also deleted in the promoter of *ZMM19* in the long-glume ‘pod corn’ due to a promoter re-arrangement (Han et al. 2012; Wingen et al. 2012; Fig 6b). Although the motifs were not highlighted in the original studies, the promoter re-arrangement (and hence the deletion of these motifs) was linked to the ectopic expression of *ZMM19* and the long-glume (*Tunicate*) phenotype. We therefore propose that the deletion of the three conserved motifs within the *SVP-A1b* promoter leads to the ectopic expression of *SVP-A1* in *T. ispahanicum* and the associated long-glume phenotype. From a mechanistic point of view, it is tempting to speculate that the absence of these motifs prevents the binding of transcriptional repressors, analogous to the proposed mechanism for *VRT-A2* (Liu et al. 2021). Consistent with this, *SVP-A1* is known to be negatively regulated by *VRN1* and *FUL2* (Li et al. 2019a), MADS-box transcription factors whose canonical binding site (CArG-box) is contained within one of the conserved motifs that is absent in the *T. ispahanicum* promoter.

### Application of larger maternal floral organ in producing plants with bigger grains

In this study, we characterized the influence of *P2* on a subset of yield components using two sets of tetraploid NIL pairs. We observed that the *P2*^*ISP*^ allele consistently increased grain length and spike length with respect to the wildtype allele in both NIL pairs (Fig 4; Supplementary Figure 8), similar to the effects seen in the F_2_ populations. However, the increase in grain length did not increase grain weight consistently, due to compensatory effects on grain width and possibly grain depth (Supplementary Figure 8). The increase in spike and grain length were equivalent to those seen in the hexaploid *P1* NILs, although *P1* significantly increased grain weight in a consistent manner. Despite these general similarities, we also detected differences between the *P2* and *P1* NILs. For *P2*, we did not detect any differences in heading time nor height (Supplementary Figure 9), unlike *P1* which delayed heading and increased plant height (Adamski et al 2021). Similarly, we observed a consistent decrease in the number of grains per spike in the *P2* NILs (due to a lower number of fertile florets per spikelet; Supplementary Figure 10), whereas we did not detect significant differences in grain number across three years in *P1* NILs. It is important to note however, that the tetraploid *P2* NILs are less adapted to UK growing conditions than the *P1* NILs, which have a UK spring wheat recurrent parent background. Further evaluation of *P2* in an equivalent UK background would be warranted to accurately assess the effect of *P2* on yield components. Despite this limitation, we observed a robust effect of *P2* on grain length and floral organ size.

There is a strong correlation between the size of the grain and that of the floral organs (Millet 1986). Here, we show that the *T. ispahanicum P2* allele increases the length of lemma and palea, in addition to grain length (Fig 4), similar to the *P1* allele of *T. polonicum* and *T. petropavlovskyi*. Several potential mechanisms have been proposed to explain this correlation. First, the floral organs (lemma/palea in wheat, referred to as hulls in rice) are proposed to physically limit grain size in cereals, so increases in floral organ size would allow more space for grains to grow into (indirect effect). In rice, knockout of *short grain6* (*OsSG6*), an AT-rich sequence and zinc-binding protein was shown to reduce hull cell division resulting in a smaller hulls and smaller grains (Zhou and Xue 2020). Since *OsSG6* was strongly expressed in the hulls, but not in the endosperm, the authors proposed that the change in grain size was due to the change in floral organ size. Additional examples in rice include *OsOTUB1, OsGW2, OsWRKY53*, and *OsCYP78A13* where hull size is modulated via either cell proliferation or cell expansion and is accompanied by a change in grain size (Huang et al. 2017; Song et al. 2007; Tian et al. 2017; Yang et al. 2013, as reviewed in Li and Li 2015; Li et al. 2019b). These studies, however, did not provide evidence that the changes in grain size were a result of changes in growth space. Alternatively, genes that influence the size of floral organs can have pleiotropic effects that can also influence grain size directly. In barley, *HvAP2* influences the size of both grains and maternal floral organs independently (Shoesmith et al. 2021). *HvAP2* represses cell expansion in maternal floral organs, but also limits both cell length and cell number in the grain pericarp tissue to reduce grain size. Similarly, in *T. polonicum*, the increase in the length of grains and maternal floral organs is accompanied by elevated/ectopic expression of *VRT-A2* in both tissues (Adamski et al. 2021). However, a mechanistic link between ectopic expression in the grain and increased grain length has not been established as in *HvAP2*.

In the case of *P2*, the differences in floral organ size are already established at anthesis, although carpel size is similar between the NILs at this stage. *SVP-A1* expression remains higher in the *P2*^*ISP*^ grains only during the first few days of grain development (non-significant differences by 10 dpa; Fig 5b), which coincides with the first observed differences in grain length between the NILs at 5 dpa (Fig 4b). At this very early stage it is unlikely that the floral organs physically constrain grain length as the developing grain is less than 50% of its final length. This supports a more direct role of *P2* on grain growth as shown for *HvAP2*. This is in contrast to *P1*, where grain length differences were first observed at 14 days post anthesis by which time grains had reached > 75% of their final length. The precise mechanisms, however, by which *P2* (and *P1*/*VRT-A2*) directly and/or indirectly affect grain length remains to be determined.

## Supplementary Table Captions

**Supplementary Table 1. Initial mapping of *P2* long glume phenotype using seven F**_**3:4**_ **homozygous recombinant lines from the T1 × LDN and TRI × LDN populations. Glume length is reported as ± standard error of the mean. The *P* value is based on a *t*-test between recombinant (Rec) and non-recombinant (Non-Rec) lines within each recombinant family**.

**Supplementary Table 2. Fine mapping of the *P2* long glume phenotype using eleven critical F**_**3:4**_ **homozygous recombinant lines from the T1 × LDN and TRI × LDN populations. Glume length is reported as ± standard error of the mean. The *P* value is based on a *t*-test between recombinant (Rec) and non-recombinant (Non-Rec) lines within each recombinant family**

**Supplementary Table 3. List of KASP markers used for mapping of *P2* in the T1×LDN, TRI×LDN, and LD222× P2-LD222 F2 populations and the results from QTL mapping**.

**Supplementary Table 4. 35K Axiom array genotyping result of the *P2* near-isogenic lines P2-LD222 (BC**_**6**_**) and its recurrent parent LD222. Conversion type refers to the quality of the marker; PolyHighRes, NoMinorHom and OTV markers were considered high quality**.

**Supplementary Table 5. Effect of *P2* haplotype group on spike length and spikelet number in the T1 × LDN and TRI × LDN populations. \**P* value based on planned contrast between haplotype groups within each population**.

**Supplementary Table 6. Summary table of the effect of *P2* on area, length and width of glume, lemma, grain and palea across different regions of the spike in LD222 NILs. The apical region contains the data from the apical 25% of the spikelets, central region contains the data from middle 50% while the basal region contains the basal 25% of the spikelets. \**P* value is based on planned contrast between *P2***^***WT***^ **and *P2***^***ISP***^ **at the specific position (black *P* value *NS*, blue *P* < 0.05, red *P* < 0.01)**.

**Supplementary Table 7. List of primer sequences for RT-qPCR (*SVP-A1, SVP-B1, VRT-A2, VRT-B2*), PCR primers and KASP assay for the 482-bp promoter deletion, and KASP markers for the SVP-A1b Q144R polymorphism**.

**Supplementary Table 8. Summary table of the germplasm screened for the presence of *SVP-A1b* (482-bp promoter deletion and Q144R polymorphism)**.

**Supplementary Table 9. Germplasm from Kuckuck’s expedition that was screened for the presence of *SVP-A1b*. Coordinates of collection sites are based on IPK gene bank passport data**.

**Supplementary Table 10. *VRT-B2* allele of *T. ispahanicum* accessions and Chinese Spring (RefSeqv1.0) based on gene model *TraesCS7B02G080300.1*. The last column shows the presence (Yes) or absence (No) of each polymorphism in the five sequenced genomes of *T. dicoccon***.

**Supplementary Table 11. Summary table of the individual one-way ANOVA on the effect of marker *6A:549538795, 7A:128917566* and *7B:105882162* on glume length in the T1 × LDN F**_**2**_ **population. Lsmeans within each marker with the same letter are not statistically different based on Tukey’s test (*P* > 0.01)**

**Supplementary Table 12. List of KASP markers for fine-mapping the *P2* locus using the F**_**3:4**_ **homozygous recombinants from T1 × LDN and TRI × LDN F**_**2**_ **populations**.

**Supplementary Table 13. Gene Ontology (GO) functional annotation and orthologs of 15 high-confidence genes identified within the fine-mapped region of *P2* (chromosome 6A: 549**,**037**,**133 - 550**,**717**,**813 bp)**.

**Supplementary Table 14. *SVP-A1* allele of *T. ispahanicum* accessions and Chinese Spring (RefSeqv1.0) based on the gene model *TraesCS6A02G313800.1*. The last column shows the presence (Yes) or absence (No) of each polymorphism in the five sequenced genomes of *T. dicoccon***.

**Supplementary Table 15. Summary table of ANOVA conducted on the effect of background, *P2* allele, block, and background and *P2* interaction on the phenotypes collected on LD222 and LD222 (*Rht-B1b*) NILs grown in the field. \**P* value based on planned contrast between *P2***^***WT***^ **and *P2***^***ISP***^ **within each background. **Thousand grain weight (TGW) was extrapolated based on grain weight of around 200 grains**.

**Supplementary Table 16. Summary table of ANOVA conducted on the effect of *P2*, days post anthesis (dpa), and their interaction on grain measurements for LD222 and LD222(*Rht-B1b*) NILs. Measurements were performed using the grain from the first floret of four central spikelets. \**P* value based on planned contrast between *P2***^***WT***^ **and *P2***^***ISP***^ **at each timepoint**.

**Supplementary Table 17. The effect of *P2*, pericarp cell position on the grain and their interaction on pericarp cell length in LD222 NILs. \**P* value based on planned contrast between *P2***^***WT***^ **and *P2***^***ISP***^ **at each position**.

## Statements & Declarations

### Author Contribution

Conceptualization: YC, NMA, CU; Data curation: YC; Formal analysis: YC, NMA; Funding acquisition: NMA, CU; Investigation: YC, YL, JZ, AT, NMA; Methodology: YC, NMA, CU; Project administration: NMA, CU; Resources: NW; Software: YC, NMA; Supervision: YC, NMA, CU; Visualization: YC; Writing-original draft: YC, NMA, CU; Writing-review & editing: YC, JZ, NMA, CU.

## Acknowledgement

We thank Dr. Fei Lu (Centre of Excellence for Plant and Microbial Science) for pre-publication access to whole genome sequencing data of *Triticum* accessions used in this study and Professor Lars Østergaard (JIC) for feedback on the work. We thank JIC Bioimaging facility and staff for their contribution in SEM imaging. We thank the JIC Field Experimentation and Horticultural Services teams for their support in glasshouse and field experiments.

## Data Availability

That F_2_ mapping dataset generated is included in the manuscript within supplementary data. The protein alignment of *SVP/StMADS11-like* genes in cereal is deposited in Dryad. NILs are deposited in JIC’s Germplasm Resource Unit.

