## Supplementary Figures for "The *T. ispahanicum* elongated glume locus *P2* maps to chromosome 6A and is associated with the ectopic expression of *SVP-A1*"

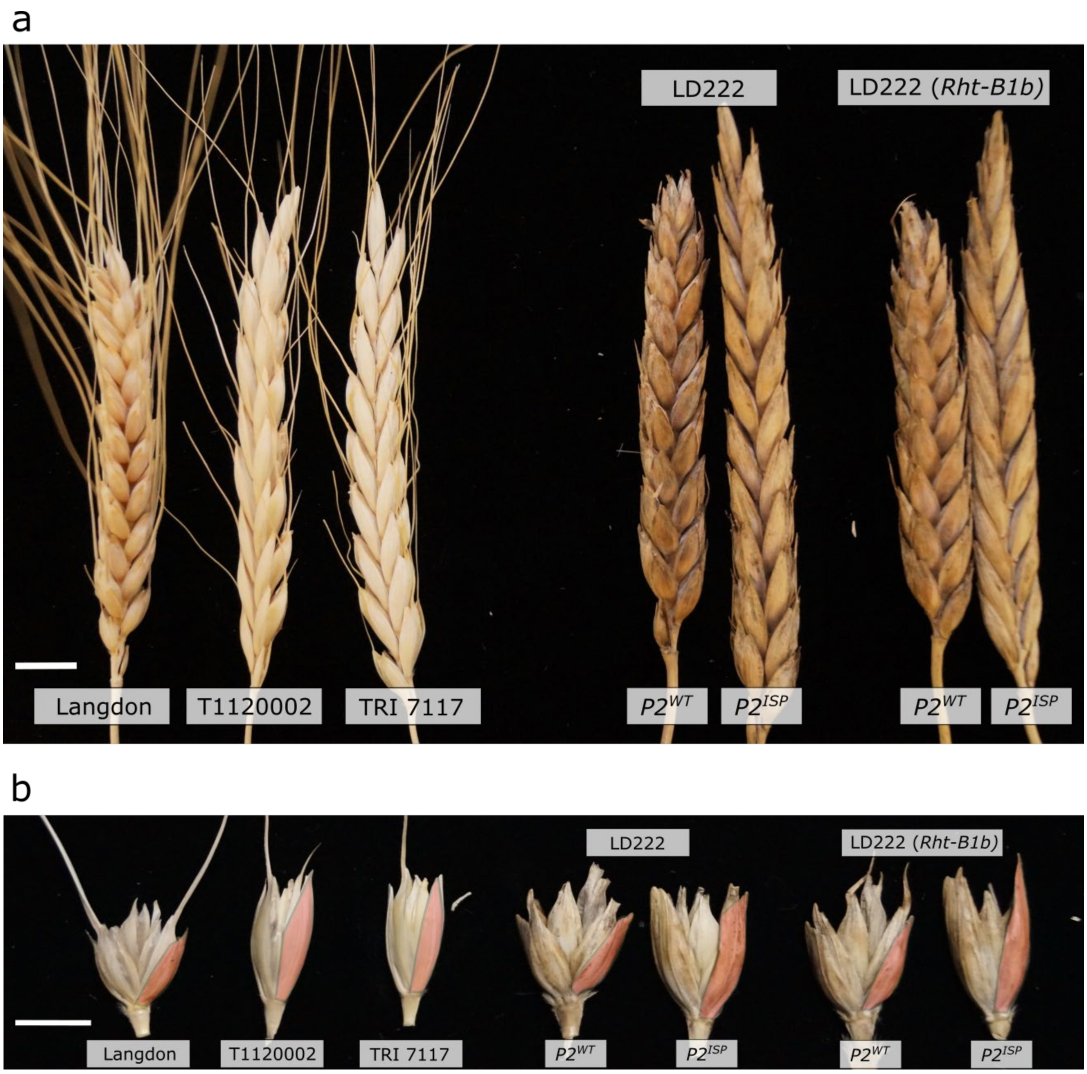


**Supplementary Figure 1. Spikes (a) and spikelet (b) of the parental tetraploid durum wheat cultivar Langdon, *T. ispahanicum* accessions T1120002 and TRI 7117, and the two sets of *P2* NILs used in the study.**

Scale bar = 1 cm. In b, glume tissues are outlined in red.

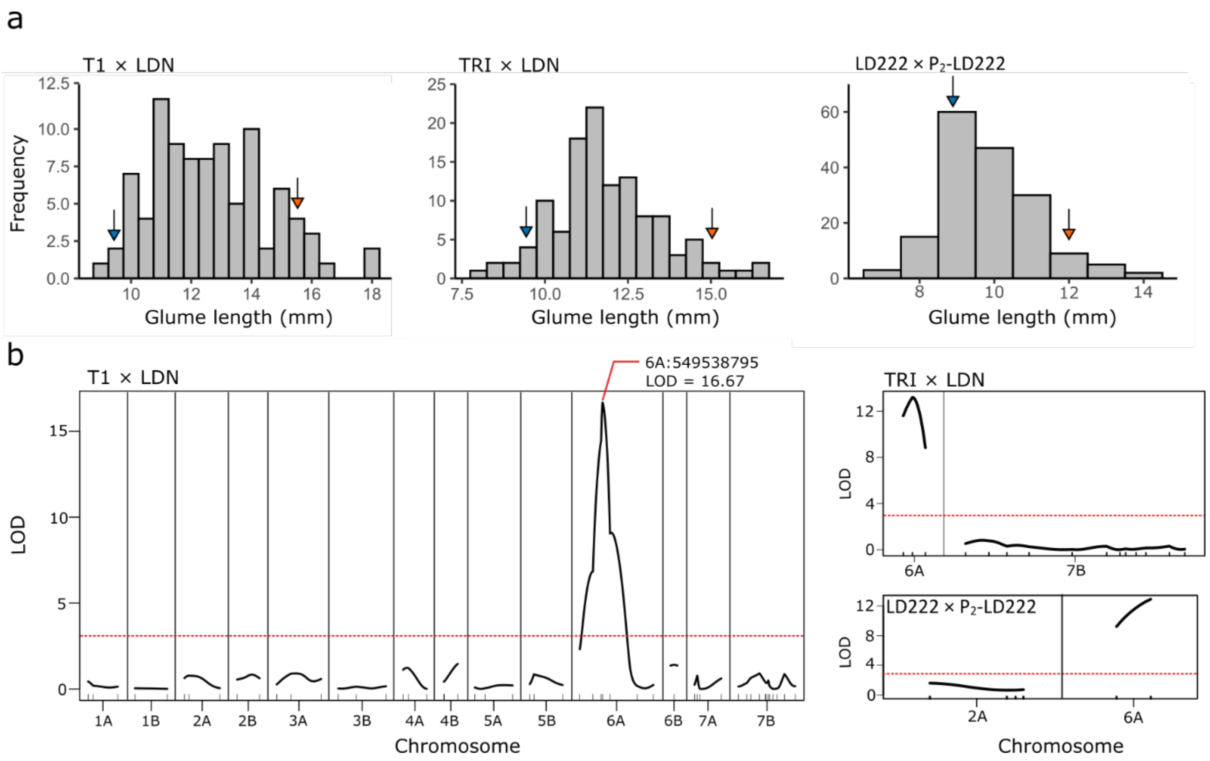


**Supplementary Figure 2. QTL analysis of glume length in three $F_2$ populations from the cross T1120002 (*T. ispahanicum*) × Langdon (T1 X LDN, *n* = 93 individuals), TRI 7117 × Langdon (TRI X LDN, *n* = 120 individuals), and LD222 × P2-LD222 (*n* = 172 individuals)**

**a**, Histograms of glume length variation in the three $F_2$ populations. Arrows indicate the glume size of $P2^{ISP}$ parent (orange) and $P2^{WT}$ parent (blue) grown in the same condition (n = 4 plants of each parental genotype). **b**, QTL plots of glume length for the three $F_2$ populations (significance threshold LOD > 3.0). The ticks represent individual markers reported in Supplementary Table 1. Markers are ordered based on genetic position. For T1 X LDN, all chromosomes have at least three markers except chromosomes 1B (2 markers), 2B (2 markers), 4B (2 markers), and 6B (1 marker).

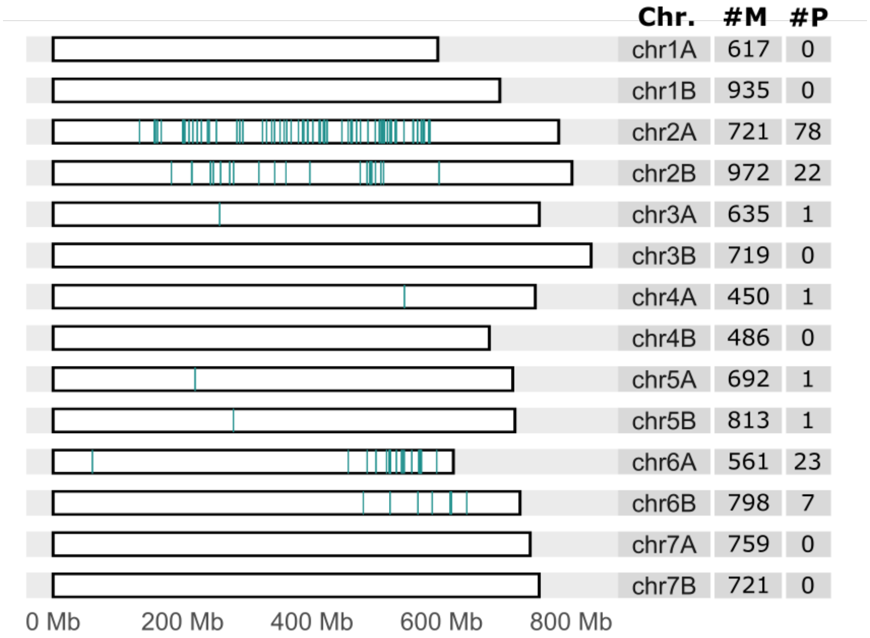


### Supplementary Figure 3. Polymorphic markers between the *P2* near-isogenic lines, LD222 and P2-LD222 using the 35K Axiom array.

The distribution of polymorphic SNPs (teal vertical lines) between the *P2* NILs across the 14 chromosomes. #M represents the number of high-quality markers after filtering (see Materials and Methods; Supplementary Table 2) on each chromosome and #P represents the number of polymorphic markers between the NILs.

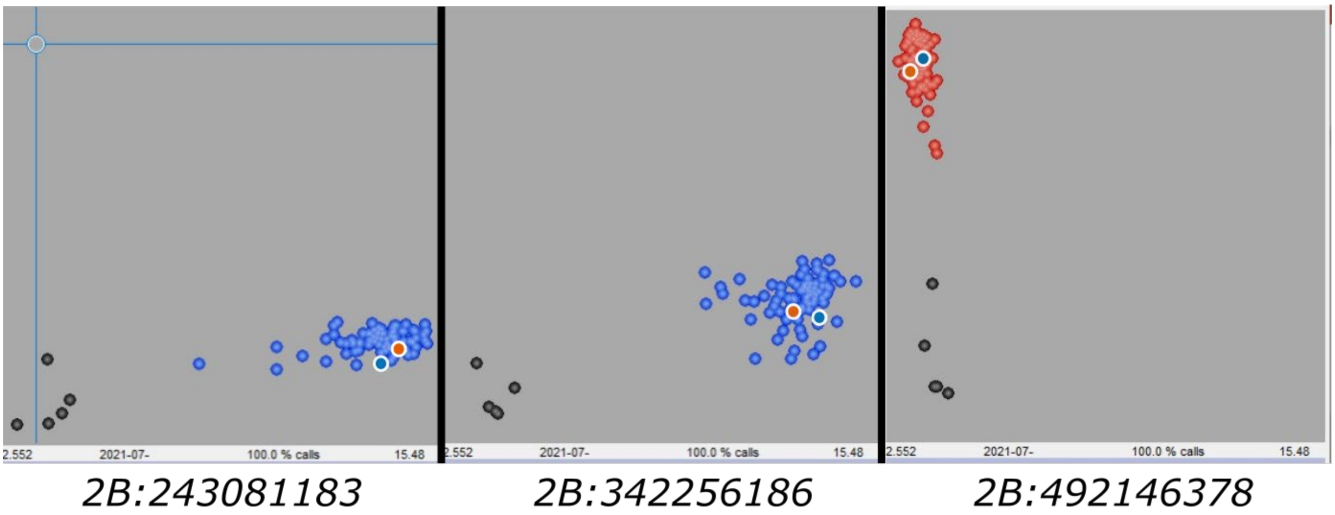


### Supplementary Figure 4. Cartesian cluster plot of the LD222 × P2-LD222 $F_2$ population genotyped with KASP markers from chromosome 2B.

X-axis is the FAM value while the Y-axis is the HEX value. Black circles are no-template controls. The parents were included in the plate as controls and are highlighted by white circles filled in with blue (LD222) and orange (P2-LD222). The 2B markers identified as polymorphic via 35K Axiom array turned out to be monomorphic in the $F_2$ population.

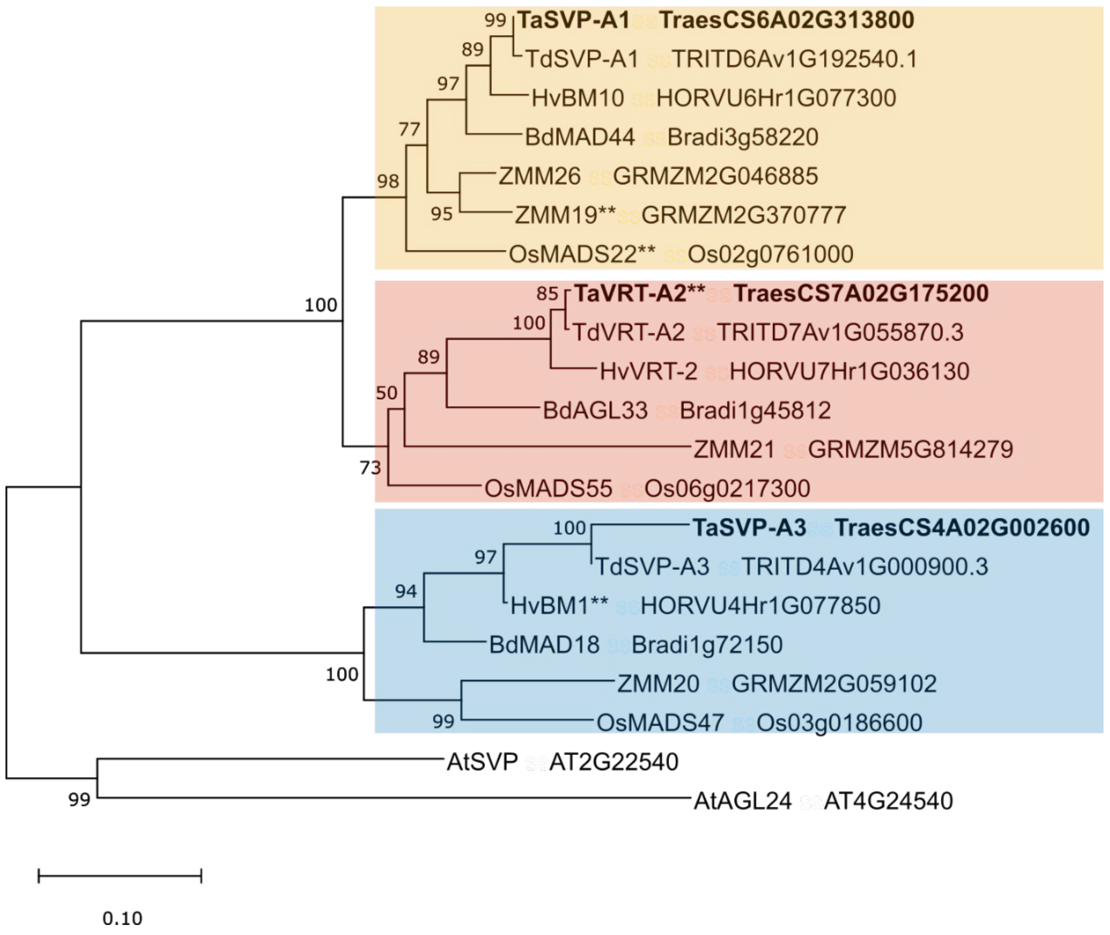


**Supplementary Figure 5. Phylogenetic tree of *SVP/StMADS11-like* proteins from *T. aestivum, T. durum, B. distachyon, H. vulgare, O. sativa,* and *Z. mays.***

The three clades of *SVP/StMADS11-like* proteins in grasses are highlighted in different background colour where the proteins from *T. aestivum* are shown in bold. ** Genes that have been shown to influence glume or lemma length (Adamski et al. 2021; Sentoku et al. 2005; Trevaskis et al. 2007; Wingen et al. 2012).

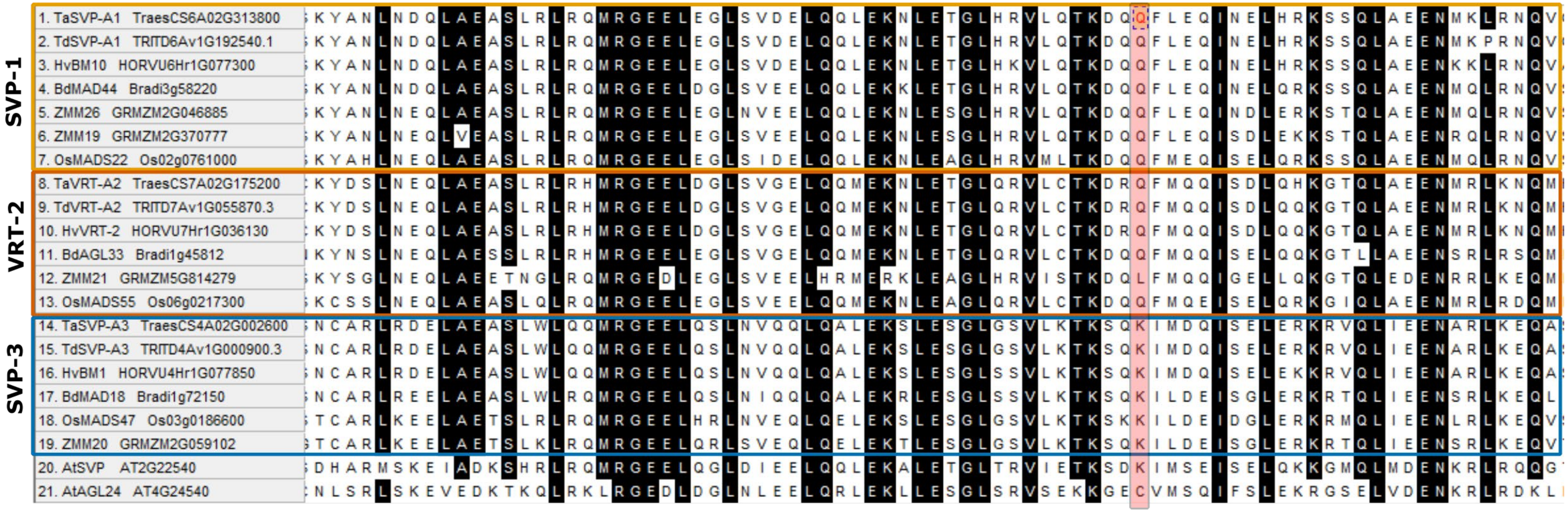


**Supplementary Figure 6. Alignment of the K-box domain (PF01486) of *SVP/StMADS11-like* proteins from *T. aestivum, T. durum*, *B. distachyon, H. vulgare, O. sativa,* and *Z. mays*.**

The alignment extends from amino acid 92 to 170 of SVP-A1. The three clades of *SVP/StMADS11-like* proteins (Supplementary Figure 5) are indicated by coloured boxes; two *Arabidopsis* SVP proteins (*AtSVP* and *AtAGL24*) are also included in the alignment. The position of the Q144R missense mutation is highlighted in red. Amino acids that are conserved across 90% of the input sequences are highlight in black.

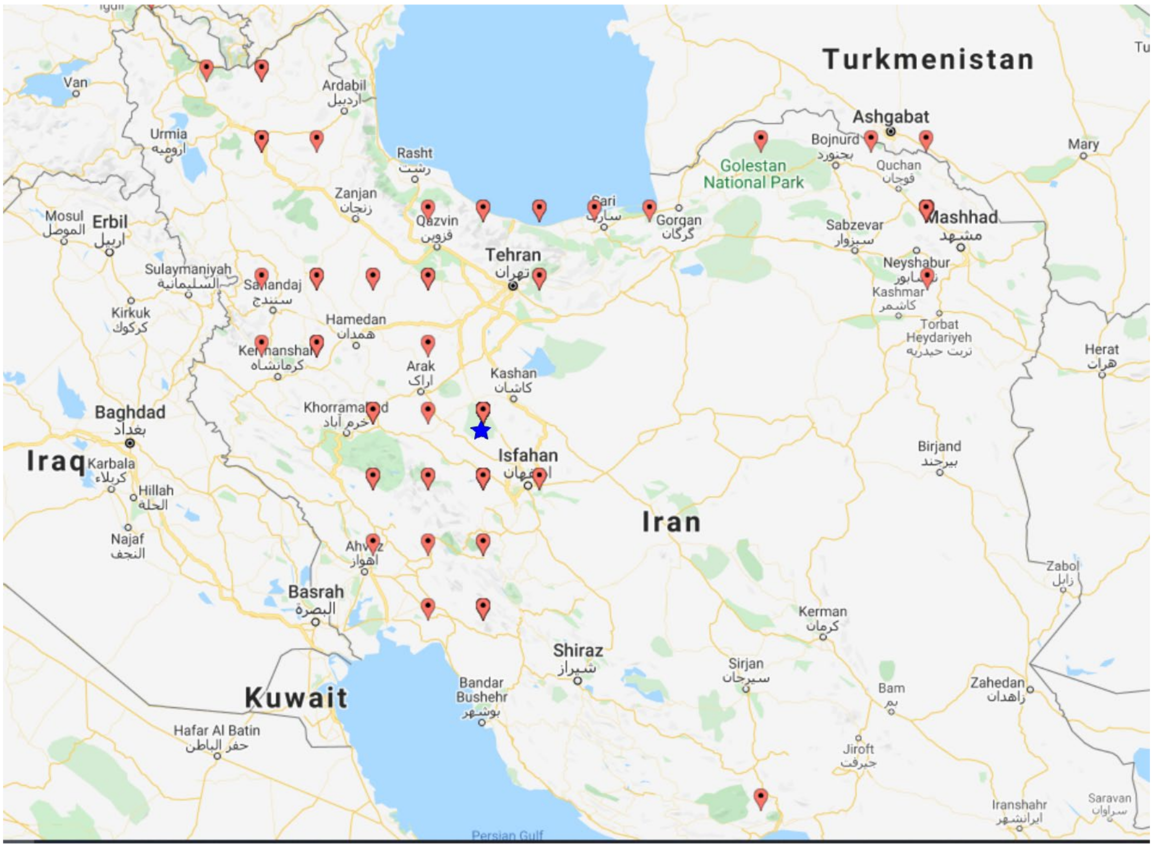


**Supplementary Figure 7. The recorded collection sites of wheat samples that were genotyped from the Kuckuck expedition (1952-1954).**

The blue star indicates where the *T. ispahanicum* accession samples were collected within the Isfahan province (Kuckuck 1956). Red flags indicate collection sites for 98 tetra- and hexaploid wheat accessions collected during the same expedition. Coordinates for these accessions are based on passport data from the IPK Genebank (Supplementary Table 9).

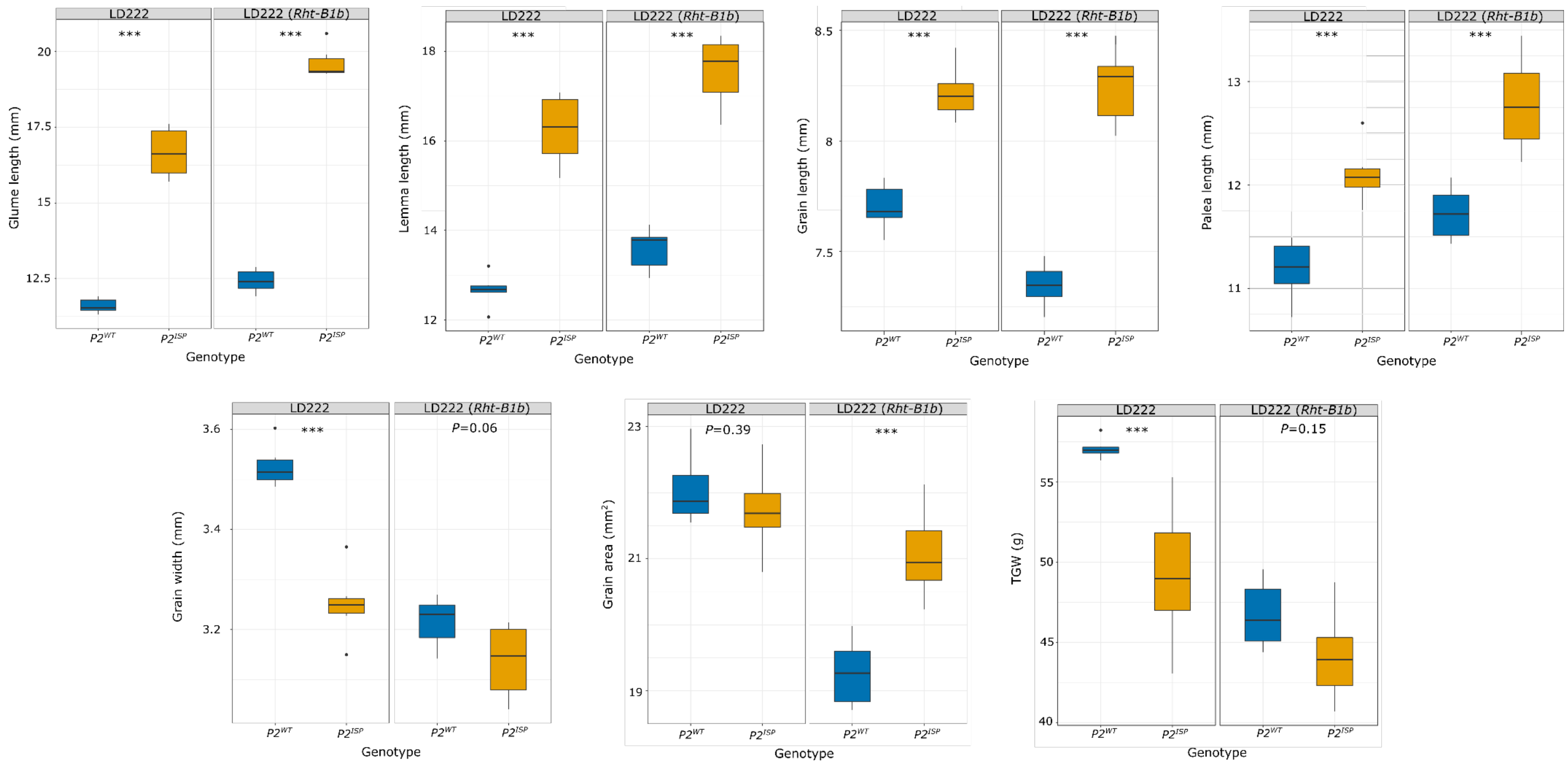


**Supplementary Figure 8. Phenotypic effect of *P2* on glume, floral organs, and grain morphology measured in LD222 and LD222(*Rht-B1b*) NILs grown in a randomized complete block design (*n* = 6 blocks) field trial.**
Measurements were performed on five representative primary spikes within each of the six biological replicates (blocks). Glume, lemma and palea length were based on the tissues from the first floret of the four central spikelets. LD222(*Rht-B1b*) NIL pair allow for the evaluation of *P2* effect under semi-dwarf background. Grain related traits were measured on all the grains from the five spikes using a MARVIN grain analyser. Thousand grain weight (TGW) was extrapolated based on grain weight of around ~200 grains. The box plots (blue for $P2^{WT}$; orange for $P2^{ISP}$) show the middle 50% of the data with the median represented by the horizontal line. Whiskers represent datapoints within 1.5 times the interquartile range while outliers are highlighted as individual dots. *P* values based on planned contrast of the phenotype performed between $P2^{WT}$ vs $P2^{ISP}$ within each background. **P* < 0.05; ****P* < 0.001.

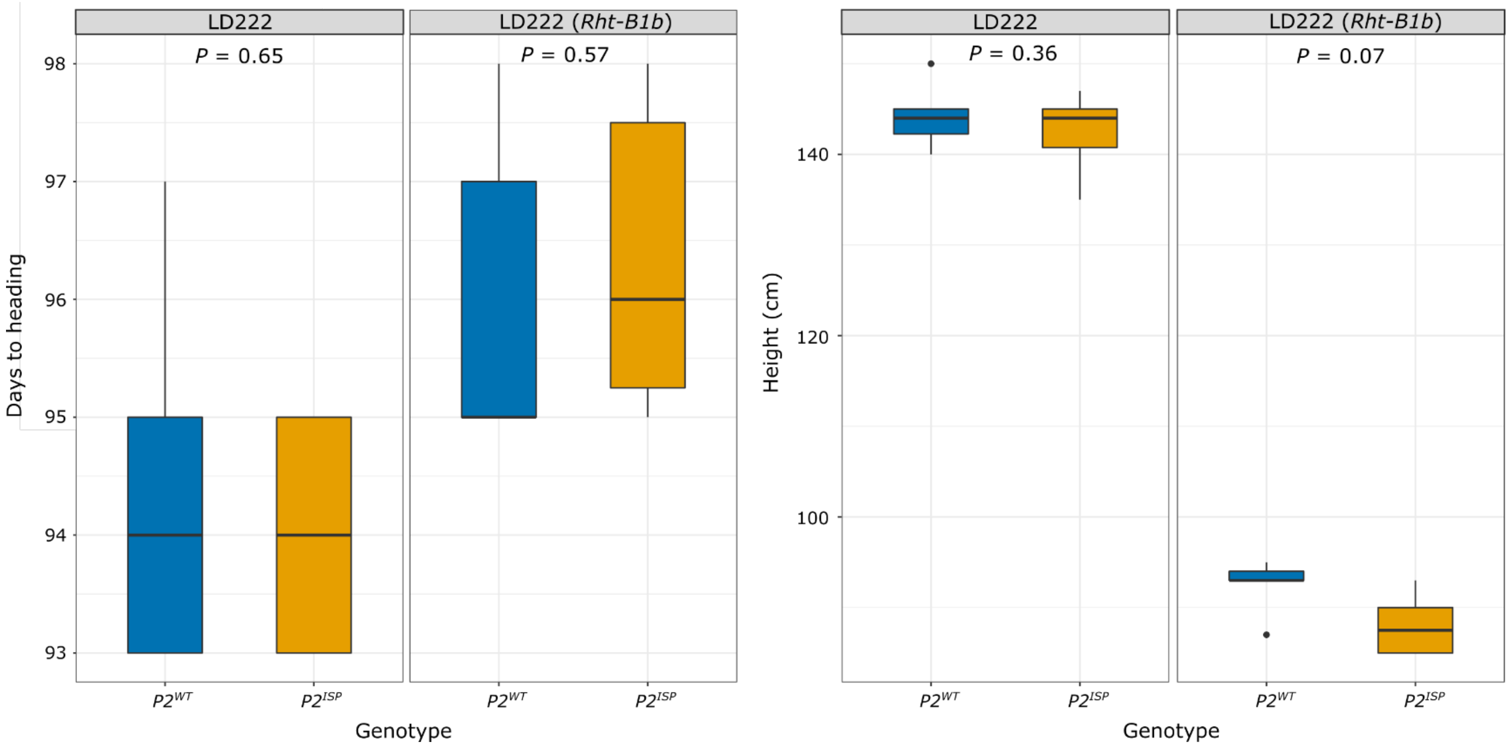


**Supplementary Figure 9. Effect of *P2* on phenology in field grown NILs in the LD222 and LD222(*Rht-B1b*) backgrounds.**

Measurements were performed on six biological replicates per genotype. LD222(*Rht-B1b*) NIL pair allow for the evaluation of *P2* effect under semi-dwarf background. The box plots (blue for $P2^{WT}$; orange for $P2^{ISP}$) show the middle 50% of the data with the median represented by the horizontal line. Whiskers represent datapoints within 1.5 times the interquartile range while outliers are highlighted as individual dots. *P* values based on planned contrast of the phenotype performed between $P2^{WT}$ vs $P2^{ISP}$ within each background.

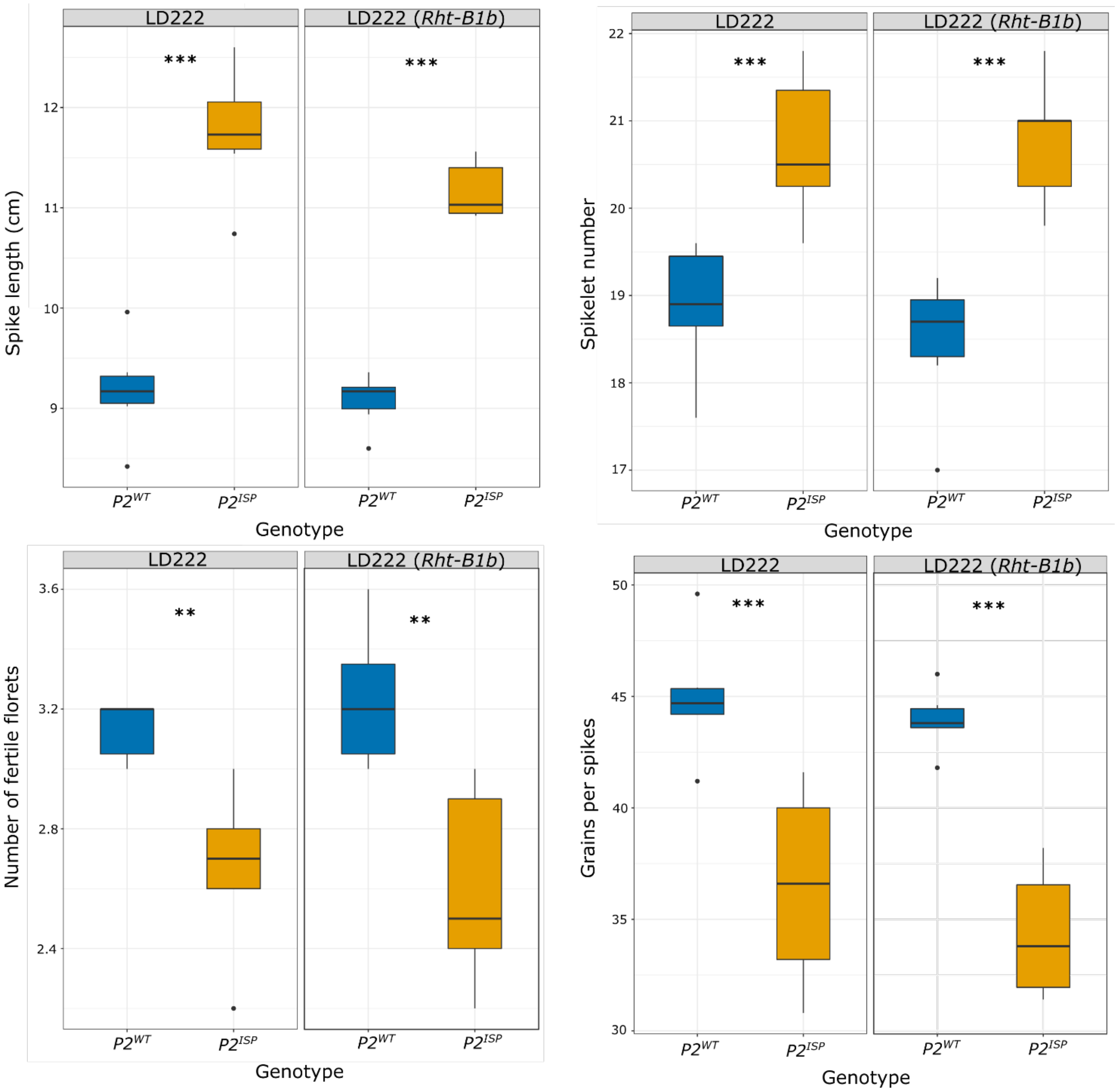


### Supplementary Figure 10. Effect of *P2* on spike morphology in field grown NILs in the LD222 and LD222(*Rht-B1b*) backgrounds.

Measurements were performed on five representative primary spikes for each of the six biological replicates per genotype. The number of fertile florets is based on the number of grains per spikelet at two central spikelets. LD222(*Rht-B1b*) NIL pair allow for the evaluation of *P2* effect under semi-dwarf background. The box plots (blue for $P2^{WT}$; orange for $P2^{ISP}$) show the middle 50% of the data with the median represented by the horizontal line. Whiskers represent datapoints within 1.5 times the interquartile range while outliers are highlighted as individual dots. *P* values based on planned contrast of the phenotype performed between $P2^{WT}$ vs $P2^{ISP}$ within each background. *$P < 0.05$; **$P < 0.01$; ***$P < 0.001$.

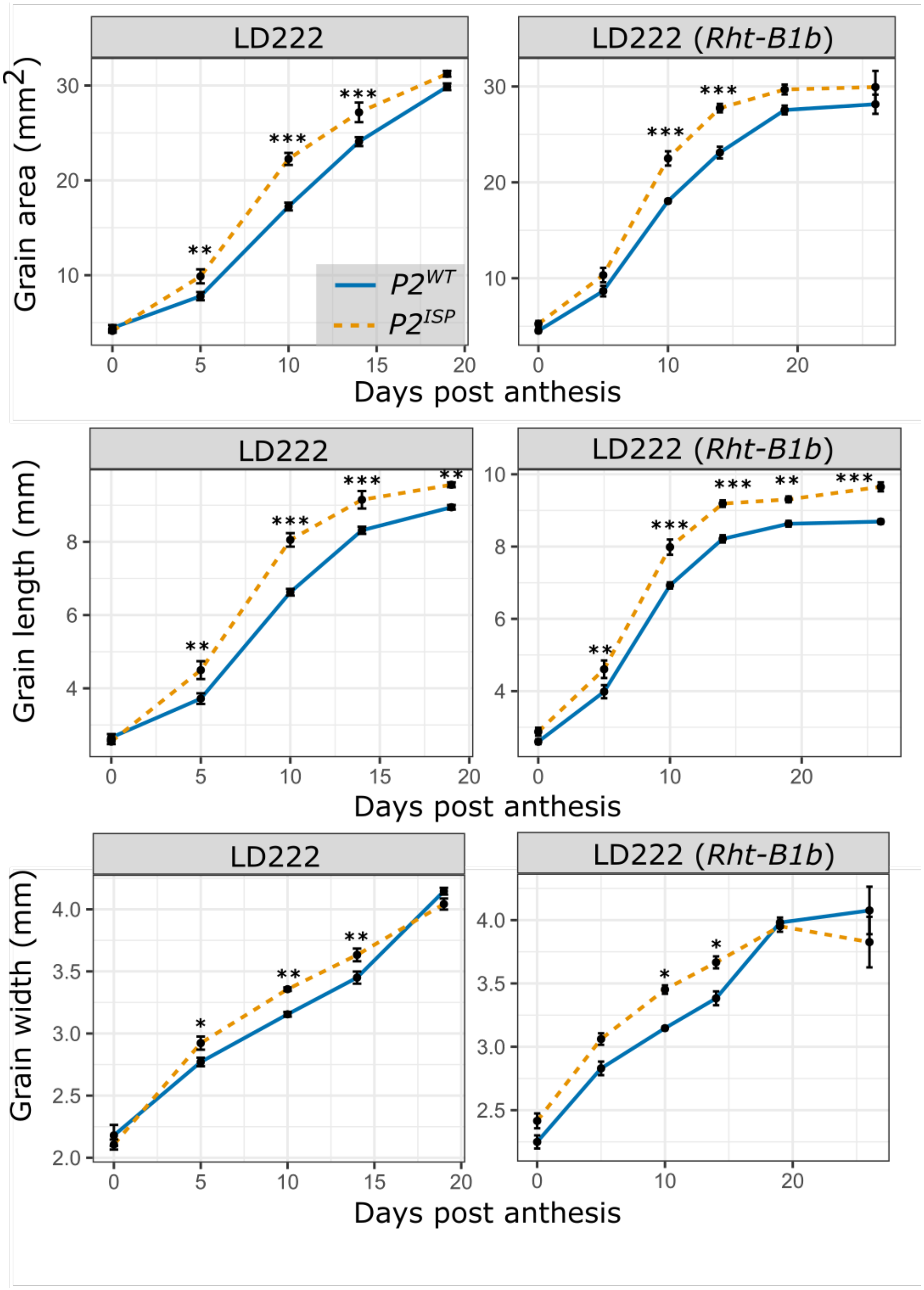


**Supplementary Figure 11. Grain development time course of field grown *P2* NILs in the LD222 and LD222(*Rht-B1b*) backgrounds.**

Measurements were performed on grains from five representative primary spikes for six biological replicates (blocks) per genotype (blue for $P2^{WT}$; orange for $P2^{ISP}$). The grains were collected from the first floret of four central spikelets for each spike. Data shows the average value of all biological replicates ± standard error of the mean. LD222(*Rht-B1b*) NIL pair allow for the evaluation of *P2* effect under semi-dwarf background. *P* values were based on planned contrast between $P2^{WT}$ and $P2^{ISP}$ within each pair of NILs at each timepoint. **P* < 0.05; ***P* < 0.01; ****P* < 0.001.
